# Starvation transforms signal encoding in *C. elegans* thermoresponsive neurons and suppresses heat avoidance via bidirectional glutamatergic and peptidergic signaling

**DOI:** 10.1101/2025.07.17.665269

**Authors:** Saurabh Thapliyal, Parvathi Sushama Gopinath, Dominique A. Glauser

**Affiliations:** Department of Biology, University of Fribourg, Chemin du Musée 10, 1700 Fribourg, Switzerland

## Abstract

Animals must continuously adapt their behavioral outputs in response to changes in internal state, including nutritional state. Here, we show that starvation induces a profound and progressive suppression of thermonociceptive behavior in *Caenorhabditis elegans*. During early food deprivation (1-hr off food), the thermoresponsive sensory neurons AWCs mediate robust heat-evoked reversals over a broad range of stimulus intensities via glutamate and FLP-6 neuropeptide signaling, each covering distinct heat intensity ranges. After six hours of food deprivation (prolonged starvation), heat-evoked reversal responses are nearly abolished, independently of any external food odor cues. Starvation triggers a shift in the distribution of AWC heat-evoked calcium response polarity, transitioning from mostly excitatory responses to a more heterogeneous pattern combining both excitatory and inhibitory activities. This switch relies on ASI neurons, proposed to work as internal state-sensing neurons. INS-32 and NLP-18 neuropeptide signals from ASI switch from a reversal-promoting to a reversal-inhibiting effect. In addition, reversal-promoting glutamatergic transmission by AWC is antagonized by glutamatergic transmission from non-AWC neurons that suppresses FLP-6-dependent reversals. Our findings define a circuit logic by which nociceptive responsiveness gating by internal nutritional state is linked to dynamic modulation of sensory neuron activity patterns and orchestrated by bidirectional glutamatergic and neuropeptidergic signals. More broadly, this study illustrates how sensory systems integrate metabolic information to prioritize behavioral outputs under changing physiological conditions, providing mechanistic insight into the plastic coupling between sensation, internal state, and action selection.

## INTRODUCTION

To ensure survival, animals must detect and avoid noxious stimuli, such as harmful chemicals, toxic gases, or extreme temperatures. These avoidance behaviors, often categorized as nociceptive, are conserved across phyla and essential for minimizing damage and guiding appropriate behavioral responses. Like most animals studied so far, the nematode *Caenorhabditis elegans*, can produce robust innate escape behaviors and modulate these responses according to external and internal regulatory cues [1-4]. Many genes involved in mammalian pain perception and plasticity (such as the Transient Receptor Potential channels, CaM kinase and Calcineurin) are also implicated in worms, underscoring the evolutionary conservation of nociceptive pathways and the broad interest of the *C. elegans* model [5-9].

*C. elegans* is a small ectotherm, whose temperature equilibrates almost instantly with its surroundings. It can grow in a 15 to 25°C range. *C. elegans* produces robust avoidance behaviors in response to noxious temperature (which can be defined as temperature above 26°C) or to fast-raising thermal stimuli even below 26°C [10]. Upon acute thermal stimulation, forward-moving worms respond to head-targeted or whole animal heating by initiating backward locomotion (reversal event) and to tail-targeted heating by accelerating their forward locomotion [11]. Several thermo-responsive neurons have been characterized (see [12] for a review). Among these neurons, AFD neurons are the best characterized primary thermosensory neurons responding to temperature changes and mediating both thermotaxis in response to innocuous thermal cues and noxious heat avoidance response [13]. FLP neurons are tonic thermosensors whose activity level continuously reflects the current temperature, and which regulate animal speed and reorientation maneuvers [14, 15]. AWC neurons are polymodal sensory neurons responding to olfactory cues and to temperature in a context-dependent manner [16-18]. The pair of AWC sensory neurons asymmetrically differentiate into AWC^ON^ and AWC^OFF^, which express specific markers and respond either similarly or differentially depending on the stimulus type [19]. Regarding AWC response to temperature change, both deterministic (stimulus-locked calcium elevations upon warming [18]), and stochastic responses, whose frequency can be modulated by warming [20], have been reported depending on experimental conditions. In addition, extremely powerful IR-laser based pulses (causing a 10°C elevation from 23 to 33°C within tens of milliseconds) were shown to trigger subtype-specific response: calcium elevation in AWC^ON^ and calcium decrease in AWC^OFF^ [21]. ASI sensory neurons have also been reported to respond to change in temperature in the innocuous range [22], but their implication in noxious-heat response is not known.

Neuromodulators, including neuropeptides and monoamines, play a central role in reconfiguring behavioral responses to sensory inputs based on context [15, 23-25]. In particular, feeding state has been shown to drive behavioral plasticity across a range of paradigms in *C. elegans*, including feeding-dependent changes in locomotion and sensory processing as a response to changes in the animal’s internal physiological, sensory and/or metabolic states [26-29]. A reduction of nociceptive sensitivity or responsiveness during prolonged starvation could in principle help worms travel across harsher environments in order to find new food sources. Consistent with this potential ecological advantage, food deprivation-dependent reduction has been shown for the response to high osmolarity and chemical repellents [12, 30]. Examples of plasticity in the thermonociceptive response of *C. elegans* are so far essentially limited to the downregulation of noxious heat avoidance in response to persistent exposure to moderately noxious heat level (28°C) or repeated heat stimulations [7, 9, 31, 32]. The impact of prolonged starvation and the contribution of neuromodulation in the context-dependent regulation of thermonociceptive behaviors is largely unknown.

Here, we investigate how feeding state modulates heat avoidance behavior in *C. elegans*. We show that prolonged starvation significantly suppresses thermonociceptive responses, a plasticity not driven by external chemosensory cues but caused by nutrient availability. We identify a critical role for the AWC sensory neurons in mediating heat-evoked escape responses, and for the ASI neuroendocrine neurons in modulating these responses under starvation. Our data show that AWC neurons employ both glutamate and the neuropeptide FLP-6 to mediate avoidance, with distinct contributions from AWC^ON^ and AWC^OFF^ subtypes. Under early food deprivation, AWC responses to heat are stereotyped and excitatory, but under prolonged starvation, these responses become variable, reflecting a shift from mostly excitatory to a mix of excitatory and inhibitory responses. This shift depends on the ASI neurons, which modulate heat-evoked response via NLP-18 and INS-32 neuropeptides. Additionally, our findings suggest that glutamatergic signaling from non-AWC sources also contribute to the suppression thermonociception following starvation. Together, our results suggest a neuromodulatory mechanism by which internal nutritional state reconfigures nociceptive behavior. Interestingly, the signaling molecules and neuromodulatory mechanisms into play are separable from the ones engaged in the regulation of thermotaxis after starvation [26, 33]. Our study demonstrates that even hard-wired escape responses are not immune to physiological context, and highlights a broader principle: survival strategies in simple animals balance external threats against internal needs through circuit-level plasticity.

## RESULTS

### Starvation downregulates thermonociceptive responses in *C. elegans*

To assess how the feeding state modulates thermonociceptive behavior in *C. elegans*, we compared responses across different durations of food deprivation (Figure 1A). Synchronized first-day adult animals were stimulated with a series of 4-s infrared pulses of increasing heating power (100, 200, 300, 400 W), causing temperature increase of +2°C, +4°C +6°C and 8°C at the surface of the plate (Figure 1A). Fed animals on food produced robust heat-evoked reversal response to heat, but they also displayed a very elevated baseline of spontaneous reversals (∼38%). A 1-hour off-food condition reduced spontaneous reversals (from ∼38% to ∼10%) and led to an attenuation of heat-evoked responses at low stimulus intensities, while responses to stronger stimuli remained comparable to those of fed animals. More prolonged food deprivation led to a striking progressive reduction in thermonociceptive responses at every heating level, with responses after 6 hours of starvation approaching baseline spontaneous reversal rates (Figure 1B and C). This suggests a robust inhibition of nociceptive behavior caused by prolonged starvation. To determine whether this attenuation was due to the absence of nutrients or chemosensory cues, we conducted similar starvation experiments in the presence of food odor, with OP50 bacteria present on the petri dish lid (Figure 1D). The reduction in thermonociceptive response persisted, indicating that the effect is driven by the internal starvation state rather than external olfactory input.

**Figure 1.**
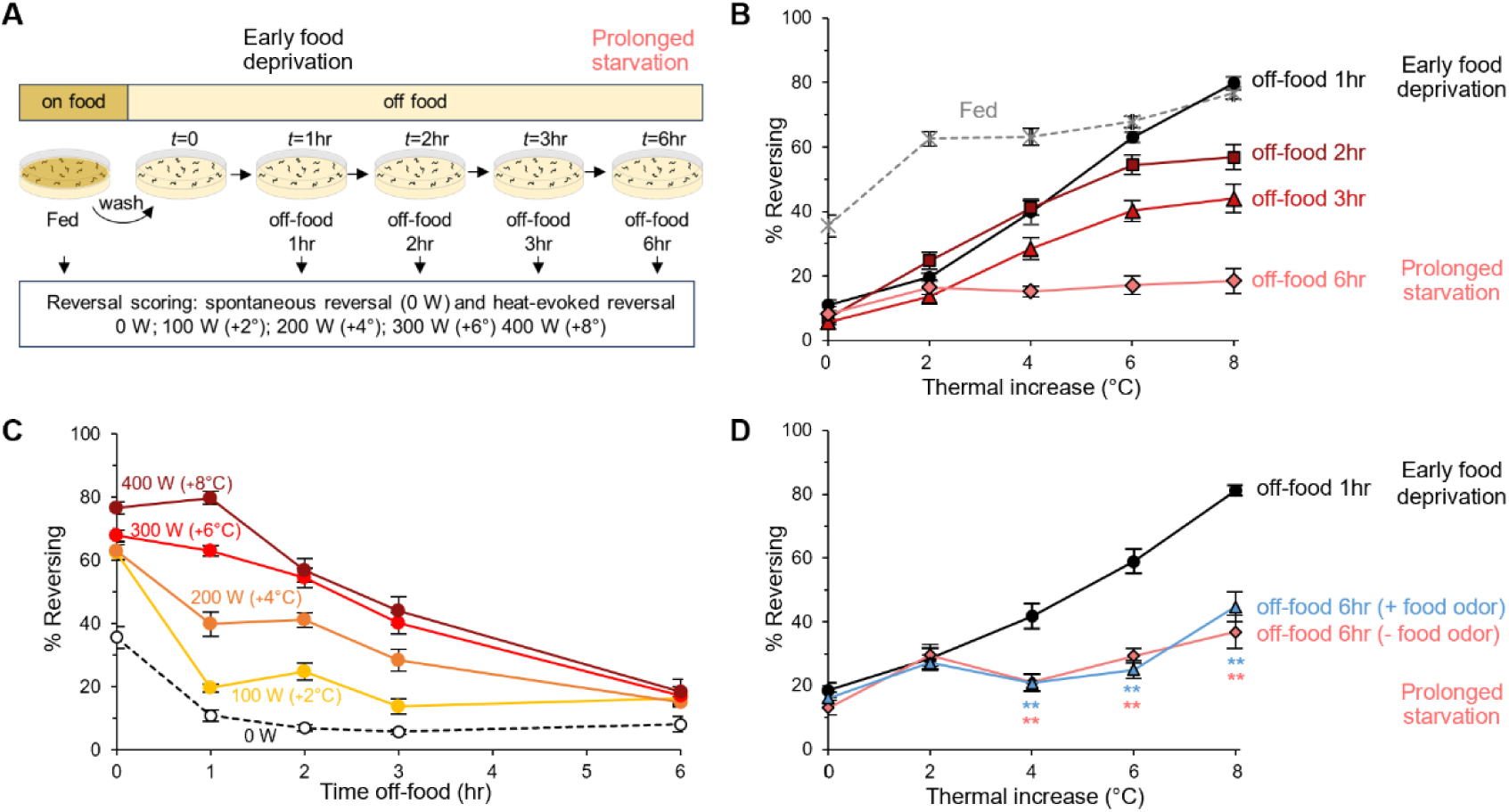
Starvation-dependent thermonociceptive plasticity in *C. elegans*. **A**. Schematic of the experimental procedure to quantify the impact of food deprivation on thermonociception in *C. elegans* adult hermaphrodites. Spontaneous and heat-evoked reversals were quantified in fed animals on food (Fed) or off food after 1, 2, 3, and 6 hr of food deprivation. Spontaneous reversals were measured as baseline reversal rate prior to any stimuli (0 W) and heat-evoked reversals were measured during a series of 4-s heat pulses at 100 W (+2°C), 200 W (+4°C), 300 W (+6°C) and 400 W (+8°C), respectively, delivered with an interstimulus interval of 20 s. **B**-**C**. Impact of food deprivation on thermonociceptive response, showing progressive attenuation of heat-evoked reversal response in the course of a 6-hr experiment. Results as average fraction of reversing animals (%) quantified in *N* ≥ 6 assays, each scoring at least 50 worms. Error bars: S.E.M. The same dataset is presented as heat dose-response curves (B) and as time-course of the heat-evoked response decrease at each heating level (C). Spontaneous reversal rate corresponds to the baseline reversal response in the absence of heating stimuli (0 W, Thermal increase = 0 °C). **D.** Effect of prolonged starvation (off-food for 6 hr) in the presence or absence of food odor during the food-deprivation period, indicating that external food odor cues cannot prevent the response reduction. **, *p>*.01 versus the early food deprivation condition (off-food 1hr), by Bonferroni post-hoc tests.

Although fed animals showed high sensitivity to noxious heat, they also displayed an elevated baseline of spontaneous reversals, which limited their utility as a control group by strongly reducing the dynamic range of heat-evoked reversal quantification and by complicating the quantitative comparison with food-deprivation conditions with much-reduced reversal baseline (Figure 1B). In addition, technical limitations in our calcium imaging setup would have prevented the intended follow-up analyses in fed animals. Based on these observations and technical considerations, we focused subsequent analyses, aiming at dissecting the circuit and molecular underpinnings of starvation-dependent plasticity, to the comparison of two off-food conditions with similar spontaneous reversal baseline: the *early food deprivation* condition (1-hour off-food, with high responsiveness to noxious heat) and the *prolonged starvation* (6-hour off-food with almost abolished noxious heat responsiveness).

### Distinct roles of AWC and ASI neurons in heat-evoked response and starvation-dependent plasticity

To identify the neural substrates underlying thermonociceptive behavior and its modulation by starvation, we genetically ablated candidate thermosensory neurons (Figure 2). Surprisingly, ablation of AFD, the canonical thermosensory neuron, had no significant effect under either 1h or 6h food deprivation (Figure 2D and Figure 2-Supplement 1). Simultaneous ablation of AFD and FLP neurons induced an increase in spontaneous reversals, but only a mild reduction in heat-evoked responses and left starvation-dependent plasticity largely intact, suggesting a minor contribution of these neurons (Figure 2-Supplement 1). By contrast, ablation of AWC or ASI neurons had dramatic effects. Removal of AWC nearly abolished heat-evoked reversal behavior across all stimulus intensities and timepoints (Figure 2B and E). While $a this observation suggests that AWC plays an essential role in mediating the thermonociceptive response under both early food deprivation and prolonged starvation. Notably, in AWC-ablated animals, the residual response level was unaffected by starvation, suggesting that AWC might also be required for the expression of starvation-dependent plasticity. In contrast, ASI ablation had almost no effect under early food deprivation but almost entirely suppressed the starvation-induced reduction in heat-evoked reversals observed after prolonged starvation (Figure 2C and E). The persistence of robust heat-evoked responses in ASI-ablated animals indicates that ASI is specifically required for starvation-dependent attenuation of thermonociceptive behavior. Together, these data suggest that, under our experimental conditions, AWC is the primary driver of heat-evoked reversals upon early food deprivation, while ASI neurons selectively mediate their suppression under prolonged starvation.

**Figure 2.**
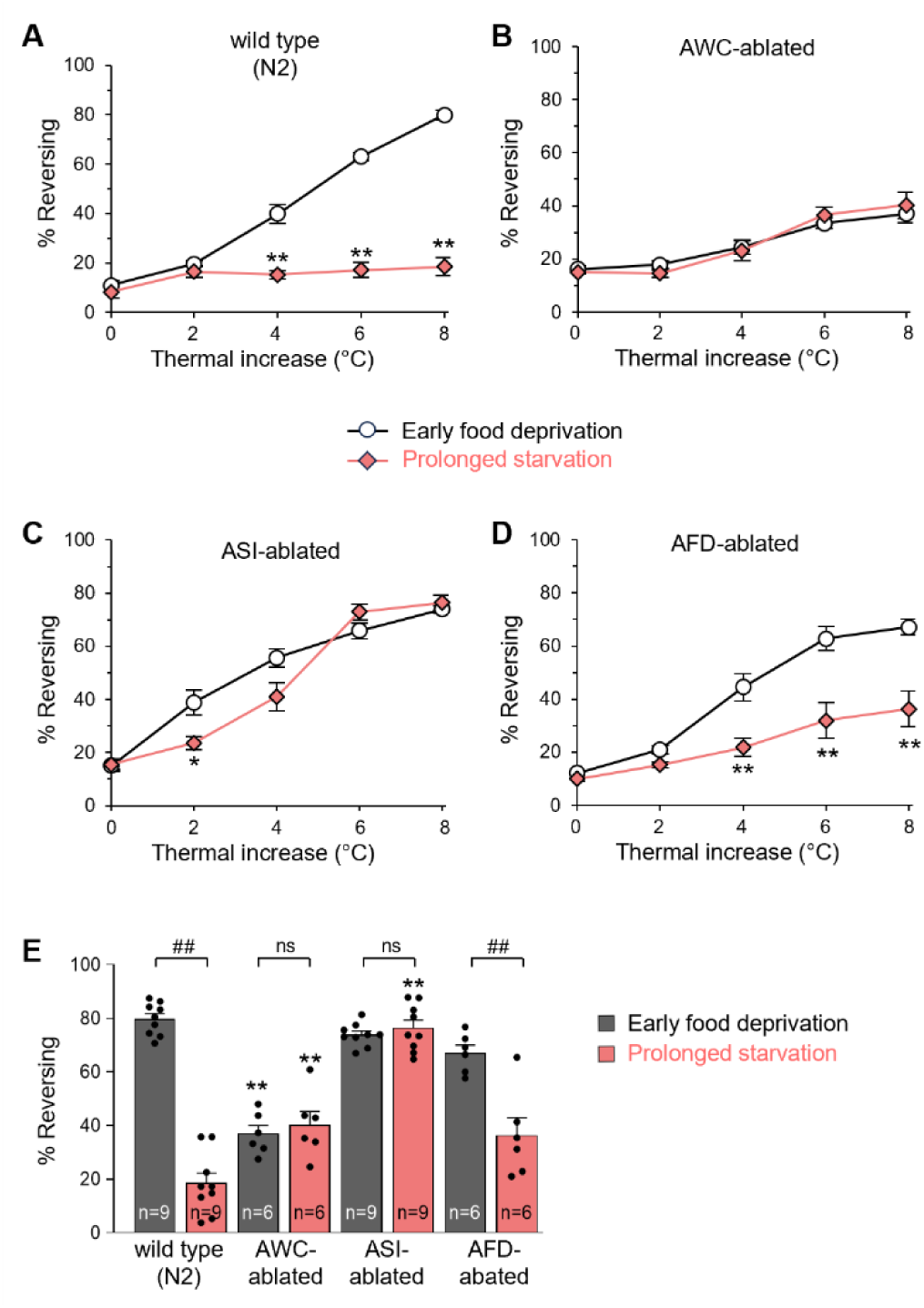
AWC mediates heat-evoked reversals upon early food deprivation and ASI mediates thermonociceptive plasticity upon starvation. Impact of the genetic ablation of AWC, ASI and AFD sensory neurons on thermonociceptive response and starvation-dependent plasticity. **A-D** Comparison of heat-evoked reversals after early food deprivation (off-food 1hr) and prolonged starvation (off-food 6 hr) in wild type and in transgenic animals with caspase-mediated ablation of indicated neurons. Results are presented as average +/- S.E.M. *, *p<.05,* and \**\*, p<.01* versus corresponding heat level in the early food deprivation condition, by Bonferroni post-hoc tests. **E** Comparison for the highest heating level (thermal increase = 8 °C) across the three genotypes presented in panel A to D. Bars as average, dots as individual assay scores, and error bars as S.E.M. *##, p<.01* between early food deprivation and prolonged starvation for each genotype; \*\**, p<.01* versus wild type (N2) at the corresponding timepoint, by Bonferroni post-hoc tests. The total number of assays (n) analyzed per condition, each scoring at least 50 worms, are indicated in panel E.

### AWC neurons mediate heat-evoked reversal via both glutamate and FLP-6 neuropeptides

We next sought to define the molecular mechanisms by which AWC neurons mediate robust heat-evoked reversals in the early food deprivation condition. AWC neurons are glutamatergic and express several neuropeptides, including FLP-6, INS-22 and NLP-5 [34, 35]. *eat-4* mutants lacking the vesicular glutamate transporter EAT-4 exhibited strong impairments in heat-evoked responses at high stimulus intensities (6 and 8°C thermal increases, Figure 3A). *flp-6* mutants displayed a marked deficit across the entire range of thermal increases (Figure 3B), while the impairment in *ins-22* and *nlp-5* mutants was less pronounced and more selectively affected the response to stimuli of intermediate intensities (Figure 3-Supplement 1). This indicates that different communication molecules are used over different thermal ranges to mediate heat-evoked reversals, which is in line with previous genetic analyses having recurrently shown that multiple genetic pathways are recruited for different levels of heat [6, 36].

**Figure 3.**
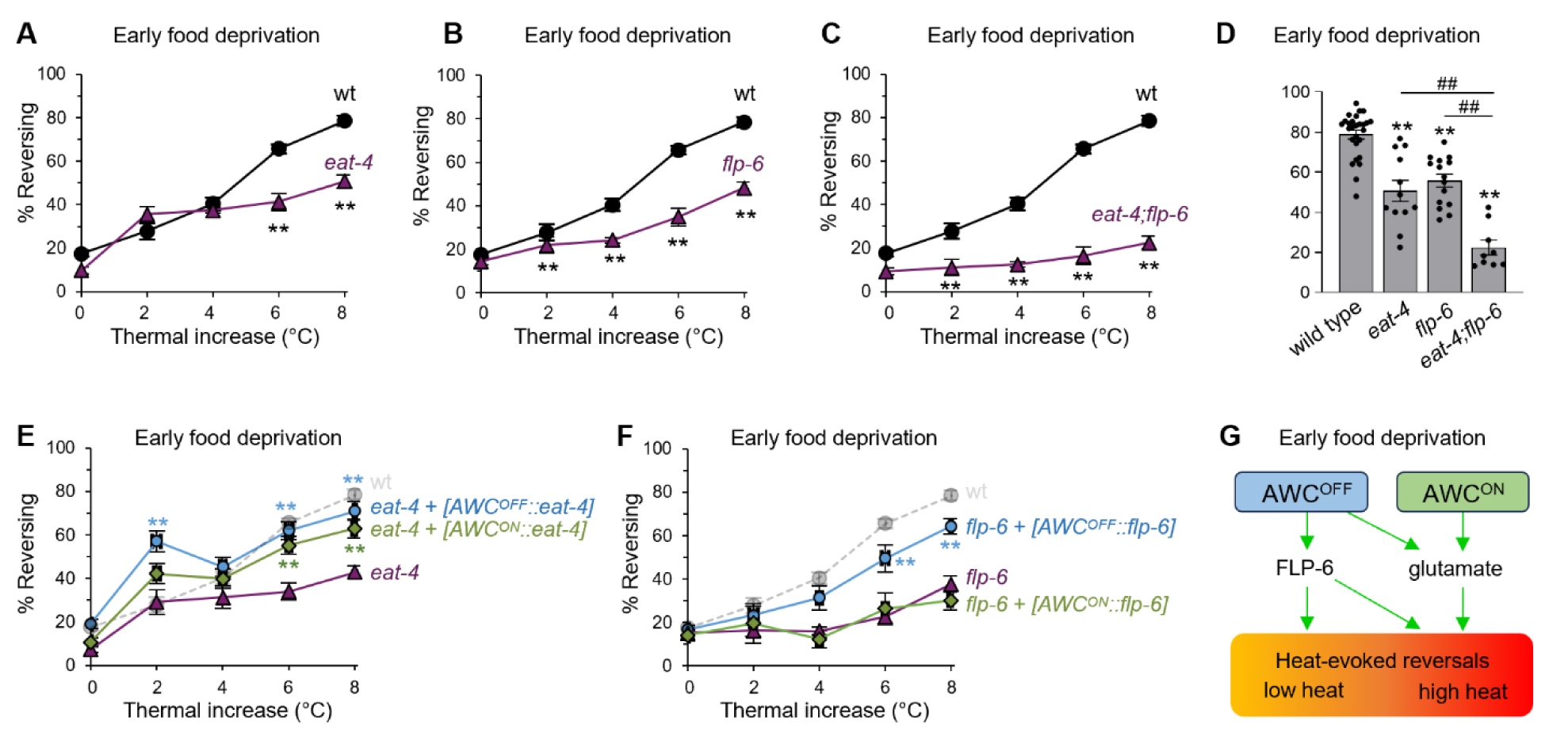
Differential engagement of glutamate and FLP-6 neuropeptides by AWC^ON^ and AWC^OFF^ in the control of spontaneous and heat-evoked reversal at various heat levels. **A, B, C, E, F** Comparison heat-evoked reversals after early food deprivation (off-food 1hr) in wild type, *eat-4(ky5)*, *flp-6(ok3056), eat-4;flp-6* double mutants and in transgenic animals with AWC subtype-specific rescue of *eat-4* and *flp-6*, respectively. Results are presented as average +/- S.E.M. \**\*, p<.01* versus wild type (A-C) and versus non-transgenic mutants (E and F) at respective thermal increase levels, by Bonferroni post-hoc tests. The number of assays (n), each scoring at least 50 worms, were : wild type, *n=*27; *eat-4*, *n=*12; *flp-6*, *n=*15; *eat-4;flp-6*, *n*=9; *eat-4+[AWC^OFF^::eat-4]*, *n=*6; *eat-4+[AWC^ON^::eat-4]*, *n=*9; *flp-6+[AWC^OFF^::flp-6]*, *n=*7; *flp-6+[AWC^OFF^::flp-6]*, *n=*8. **D** Epistasis analysis of *eat-4* and *flp-6* mutation effects on the heat-evoked reversal (thermal increase = 8 °C). Bars as average, dots as individual assay scores, and error bars as S.E.M. A two-way ANOVA showed no significant interaction of the two mutations and Bonferroni post-hoc tests confirmed significant cumulative effects of the two mutations (##, *p*<.001). **G** Visual model illustrating the specific contribution of AWC^OFF^ and AWC^ON^ to the regulation of heat-evoked reversals via the distributed action of glutamate and FLP-6 neuropeptide. A single wt control dataset is reported across panels.

Next, we focused on *eat-4* and *flp-6* mutants, showing the strongest phenotype. We addressed whether glutamate and FLP-6 signaling act dependently of each other in controlling heat-evoked reversal, by testing *eat-4;flp-6* double mutants. The residual response seen in each single mutant (Figure 3 A and B) was almost entirely abolished in the double mutant (Figure 3C). A two-way ANOVA for the highest heat stimuli with *eat-4* and *flp-6* genotypes as factors (two levels each: mutant or wild type) showed significant main effects of *eat-4* (*F_(1,67)_* =48.70, *p<*.001*, η²p*=0.421) and *flp-6* (*F_(1,67)_* =58.50, *p<*.001*, η²p*=0.466), respectively, but no interaction effects (*F_(1,67)_* =0.093, *p=*.761*, η²p*=0.001). The significant cumulative effect of the two mutations indicates that the two signaling pathways act mostly independently of each other to mediate heat-evoked reversals.

Next, we carried out cell-specific rescue to demonstrate that AWC neurons represent a relevant source of glutamate and FLP-6 neuropeptides promoting heat-evoked reversals upon early food deprivation (off-food 1hr). Transgenic rescue experiments using either *[AWC^OFF^::eat-4]* or *[AWC^ON^::eat-4]* extrachromosomal array containing-lines showed that restoring *eat-4* expression in either AWC^ON^ or AWC^OFF^ alone was sufficient to recover heat-evoked responses to near wild-type levels (Figure 3E). Interestingly, *[AWC^OFF^::eat-4]* transgene not only restored responsiveness but also enhanced reversals for low heating (+2°C) stimuli, potentially reflecting overexpression effects or altered signaling balance with other glutamatergic neurons.

Additionally, spontaneous reversal rates were restored only by *[AWC^OFF^::eat-4]*. A similar rescue approach for *flp-6* expression in AWC revealed that only *[AWC^OFF^::flp-6]*, not *[AWC^ON^::flp-6]*, could rescue heat-evoked behavior, suggesting asymmetry in neuropeptidergic signaling (Figure 3F).

Our data indicate that any one of the two AWCs might be sufficient to mediate a large part of the glutamate-dependent reversal response, but that AWC^OFF^ and AWC^ON^ might display some functional asymmetries, regarding the role of neuropeptides and the regulation of spontaneous reversal. To further address the role of AWC^ON/OFF^ asymmetry, we examined *nsy-1* mutants, developing with two AWC^ON^ and *nsy-7*, developing with two AWC^OFF^ [37, 38]. Both mutants exhibited reduced responses at high heat intensities, with *nsy-1* also showing increased spontaneous reversals (Figure 3-Supplement 2).

These results indicate that AWC neurons mediate thermonociceptive responses through both glutamatergic and FLP-6-dependent pathways, each covering distinct heat intensity ranges (Figure 3G). While either AWC subtype is sufficient for glutamatergic signaling, FLP-6 neuropeptide function appears more dependent on AWC^OFF^.

### Starvation alters the distribution of the AWC heat response polarity

To investigate if and how starvation modulates AWC activity at the cellular level, we recorded heat-evoked calcium dynamics in AWC^ON^ and AWC^OFF^ neurons using YC2.3 cameleon-expressing transgenic lines. We used a microfluidic setup to deliver defined thermal stimuli. From a baseline at 20°C, four stimuli reaching 22, 24, 26 and 28°C, respectively, were sequentially delivered, corresponding to the temperatures reached in the heat-evoked behavioral analysis (Figure 3, +2, +4, +6, +8°C, respectively). We examined responses under both early food deprivation and prolonged starvation conditions. In Figure 4A and B, we report average traces and individual traces as heat maps. Individual traces are essential to visualize the polarity of the stimulus-locked responses on a trial-by-trial and animal-by-animal basis. Globally, we noted a graded response of both AWC types, with the lowest stimulus (2 °C) causing either no response or comparatively weak response, but the higher stimuli (4, 6 and 8°C) causing obvious responses. We clustered traces as “calcium up” (traces showing mostly up regulation), “variable” (traced showing a mix of up and down regulation) and ‘calcium down’ (traces showing mostly down-regulation). The “calcium down” trials were often, but not always, followed by a “rebound” upregulation.

**Figure 4.**
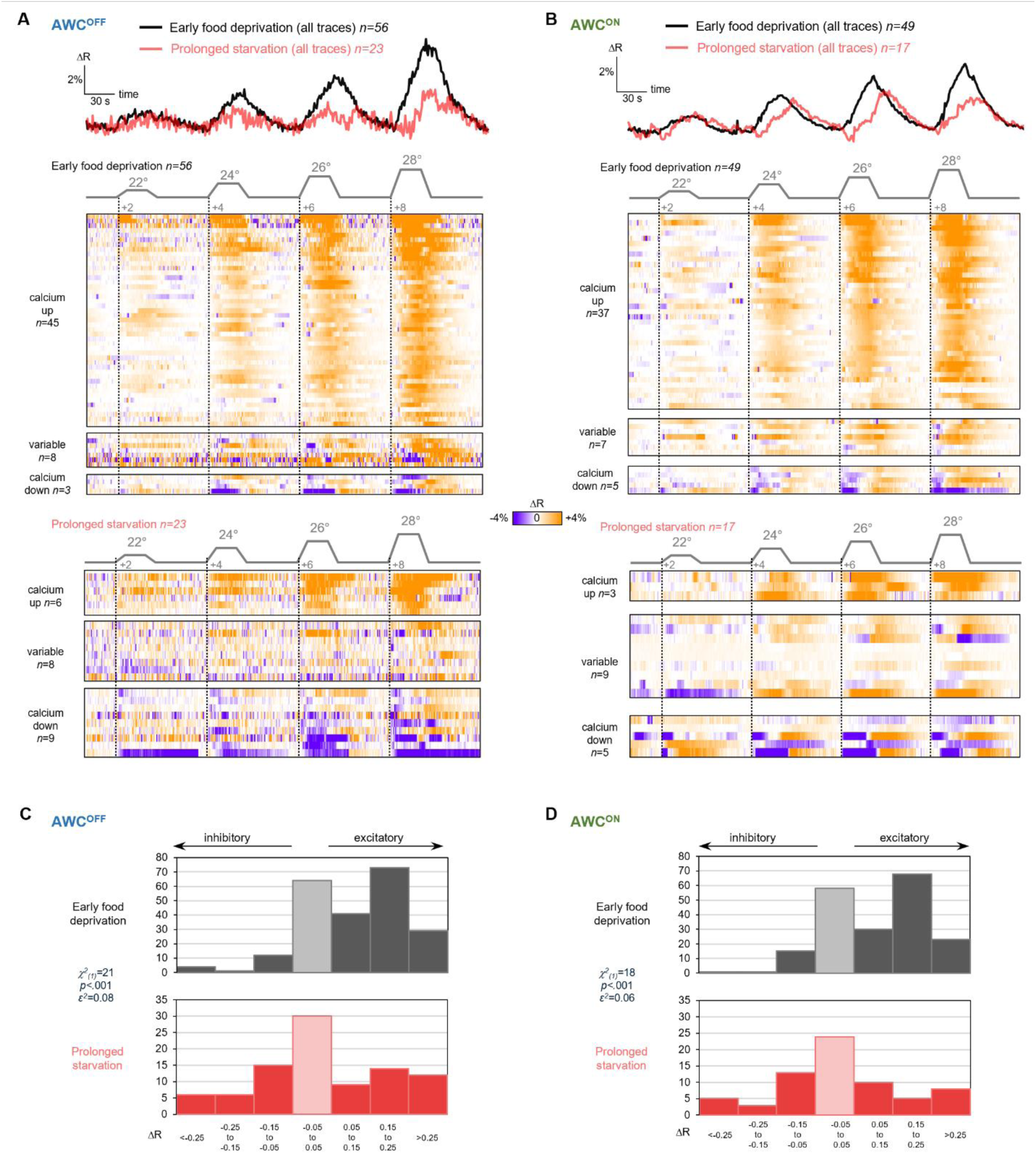
Starvation reconfigures AWC heat-evoked response polarity distribution from mostly excitatory to an heterogenous mix combining excitatory and inhibitory responses. **A-B** Calcium activity in AWC^OFF^ (left) and AWC^ON^ (right) in response to a series of four thermal up-steps. Upper plots show the average traces, lower heat maps show individual neuron traces, in the early food deprivation (off-food 1hr) and prolonged starved (off-food 6hr) conditions. **C-D** Histograms showing the distribution of calcium peak magnitudes (all thermal increase levels pooled) and highlighting the shift from mostly excitatory responses (early food deprivation, upper plot) to an equal mix of excitatory and inhibitory responses (prolonged starvation, lower plot) taking place for both AWC^ON^ and AWC^OFF^. Results of Kruskal-Wallis non-parametric tests comparing the two food-deprivation timepoints are indicated to the left of each graph.

Upon early food-deprivation, both AWC neuron subtypes responded with a stimulus-locked intensity-dependent calcium increase ("calcium up") in most animals (45/56, 80% for AWC^OFF^ and 37/49, 77% for AWC^ON^). The remaining traces showed either mixed ("variable") or "calcium down" profiles. The response peak histograms for both neuron subtypes show asymmetrical distributions with mostly excitatory responses (Figure 4C and D, upper histogram). After prolonged starvation, this response pattern shifted dramatically: the majority of traces became variable (i.e., including both calcium up and calcium down response along the stimulus series) or showed consistent calcium down-regulation (17/23, 74% for AWC^OFF^ and 14/17, 82% for AWC^ON^, both changes being statistically significant by Fisher’s exact test: *p*=1.3E-5 and *p=*7.6E-5, respectively). The response peak histograms for both neuron subtypes (Figure 4C and D) now show much more symmetrical distributions, with equally frequent excitatory and inhibitory responses. Kruskal-Wallis tests showed significant distribution differences between early food deprivation and prolonged starvation (*p<*.001, for each neuron subtype).

The starvation-dependent response shift was reflected in the average traces (Figure 4A and B) as a decreased peak amplitude and rightward shift. But the modification of the average trace was largely caused by the redistribution in response types: the dynamic of individual *calcium up* and *calcium down* traces remaining similar in each condition (Figure 4, Supplement 1 A and B). Therefore, AWC neurons, which under early food deprivation produce very consistently excitatory responses to heat (calcium increases), enter a heterogeneous and less predictable response mode under prolonged starvation, where stimuli can trigger either excitatory or inhibitory responses.

In summary, starvation does not simply attenuate stimulus-locked calcium responses in AWC; it fundamentally alters their polarity distribution, shifting from a largely predictable excitation profile to a heterogeneous regime combining excitation and inhibition. We speculate that this transformation in thermal encoding might underlie, at least in part, the behavioral plasticity observed during starvation.

### ASI is required to shift the distribution of the AWC heat response polarities

Because ASI neurons are required for the starvation-dependent thermonociceptive plasticity (Figure 2C), dispensable for noxious heat-evoked reversal response (Figure 2C), and have been previously shown to signal the feeding state of the animal [39], we hypothesized that ASI neurons might play a modulatory role to mediate the starvation impact on AWC activity patterns. To test this hypothesis, we recorded AWC^ON^ and AWC^OFF^ activity in ASI-ablated animals. In the early food-deprivation condition, 1hr off-food, ASI ablation produced no noticeable difference in AWC calcium activities (compare Figure 4 with 5). Strikingly different from the situation in animals with intact ASI (Figure 4), starvation after 6hr off-food produced no noticeable effect in ASI-ablated animals, neither for AWC^ON^, nor for AWC^OFF^ (Figure 5). In particular, the response polarity distributions remained asymmetrical with a very large majority of excitatory responses (Figure 5C and D). We conclude that the starvation-dependent shift in the distribution of AWC heat response polarity is mediated by the ASI neuromodulatory neurons.

**Figure 5.**
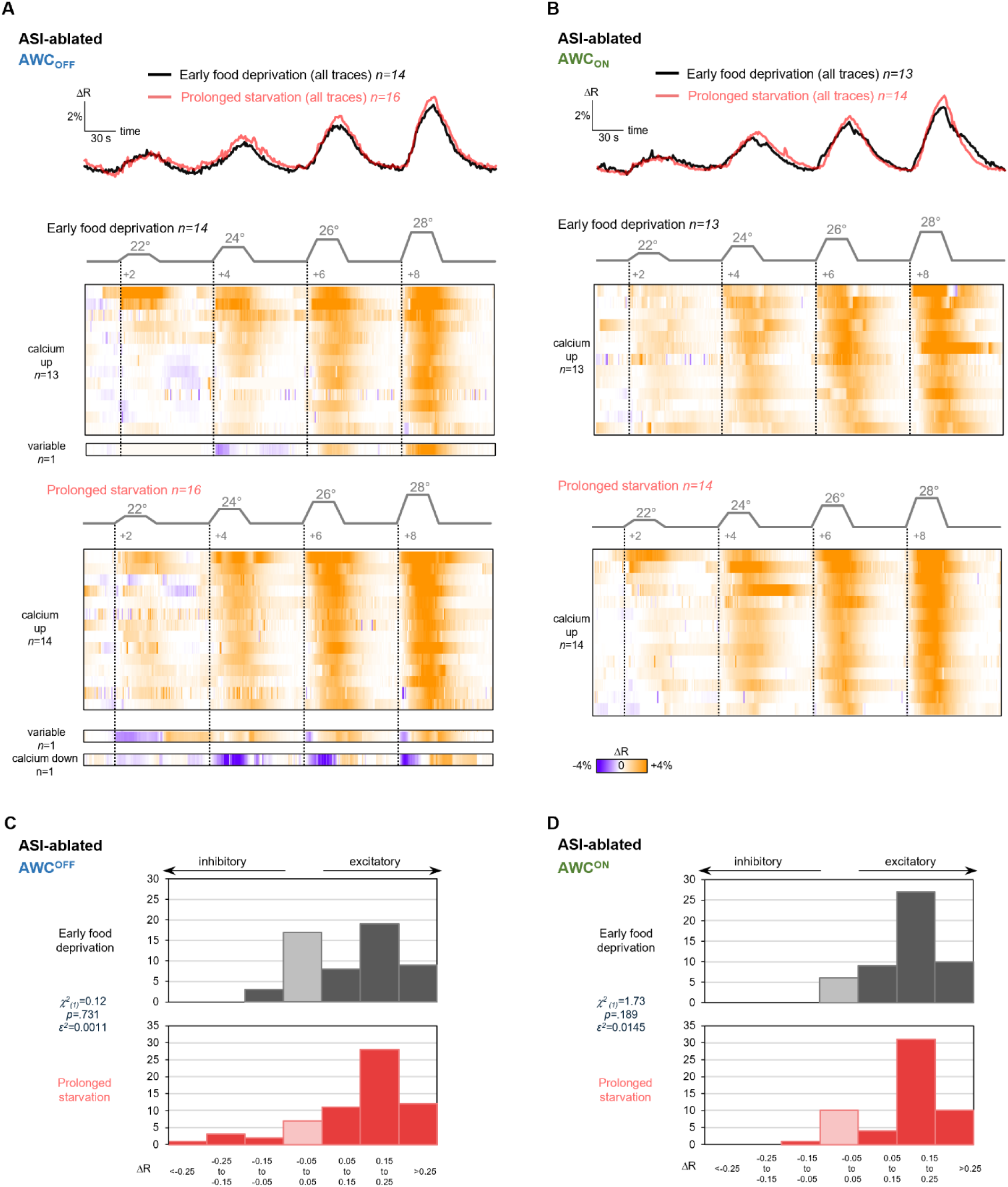
ASI ablation prevents AWC activity pattern reconfiguration upon starvation. (**A-B**) Calcium activity in AWC^OFF^ (A) and AWC^ON^ (B) in response to a series of four thermal up-steps in ASI-ablated animals. Upper plots show the average traces, lower heat maps show individual neuron traces, in the early food deprivation (off-food 1hr) and prolonged starved (off-food 6hr) conditions. The starvation impact seen in animals with intact ASI (Figure 4) is absent when ASI is ablated. **C-D** Histograms showing the distribution of calcium peak magnitudes (all thermal increase levels pooled) for both AWC^ON^ and AWC^OFF^. Results of Kruskal-Wallis non-parametric tests comparing the two food-deprivation timepoints are indicated to the left of each graph.

### Starvation-evoked thermonociceptivce plasticity is promoted by ASI-expressed neuropeptides, including NLP-18 and INS-32, and modulated by glutamatergic signaling

To elucidate the molecular signaling pathway underlying starvation-evoked thermonociceptive plasticity involving the AWC and ASI sensory neurons, we investigated the role of candidate neurotransmitters, including INS-1 which was previously shown to modulate thermotaxis in response to starvation [26], as well as other neurotransmitter/neuropeptides expressed in AWC and ASI.

First, we examined the phenotype of *ins-1* mutants, defective for starvation-evoked thermotaxis plasticity [26]. Interestingly, the thermonociceptive plasticity in these mutants was intact (Figure 6-Supplement 1), indicating that the molecular signaling controlling thermo-nociceptive plasticity is at least in part different from that engaged for negative thermotaxis plasticity as previously studied [26].

Second, we tested whether starvation-dependent plasticity was preserved in *eat-4* and *flp-6* mutant backgrounds, which we had suggested to represent the main AWC transmitters controlling heat-evoked reversals under the early food deprivation condition (Figure 3). Even if the heat-evoked response upon early food-deprivation was reduced relative to wild type in *flp-6* mutants, a significant further decline was seen after prolonged starvation (Figure 6B). In contrast, *eat-4* mutants displayed markedly elevated heat-evoked responses after prolonged starvation, even exceeding the response level seen in the early food deprivation condition for low heat stimuli (Figure 6C). This potentiated response in *eat-4* mutants was entirely dependent on an intact FLP-6 signaling, since reversal responses in *eat-4; flp-6* mutants were entirely abolished, like in *flp-6* single mutant (Figure 6C-E, a two-way ANOVA indicating a significant interaction effect between the two mutations: *F_(1,70)_* =0.093, *p<*.001*, η²p*=0.247). Moreover, the potentiated response in *eat-4* single mutant could not be rescued by expressing *eat-4* rescue transgene selectively in either AWC^OFF^ or AWC^ON^ neurons (Figure 6F). Interestingly, AWC^OFF^-specific rescue produced a further potentiation of heat-evoked reversal response to high heat stimuli (Figure 6F, 6 and 8°C thermal increases), aggravating the phenotype of *eat-4* mutants. These results are consistent with a model in which glutamatergic signaling regulates heat-evoked reversals in starved animals via two bidirectional drives (Figure 6G). On the one hand, glutamatergic signaling—originating from AWC^OFF^—up-regulates reversals in response to high heat stimuli, thus contributing to prevent starvation-induced thermonociceptive plasticity. On the other hand, glutamatergic signaling—originating from neurons other than AWC— down-regulates reversals over a broad range of heat intensities, thus promoting starvation-induced thermonociceptive plasticity. The latter glutamatergic signaling inhibitory effect seems to be more dominant and to depend on intact FLP-6 signaling.

**Figure 6.**
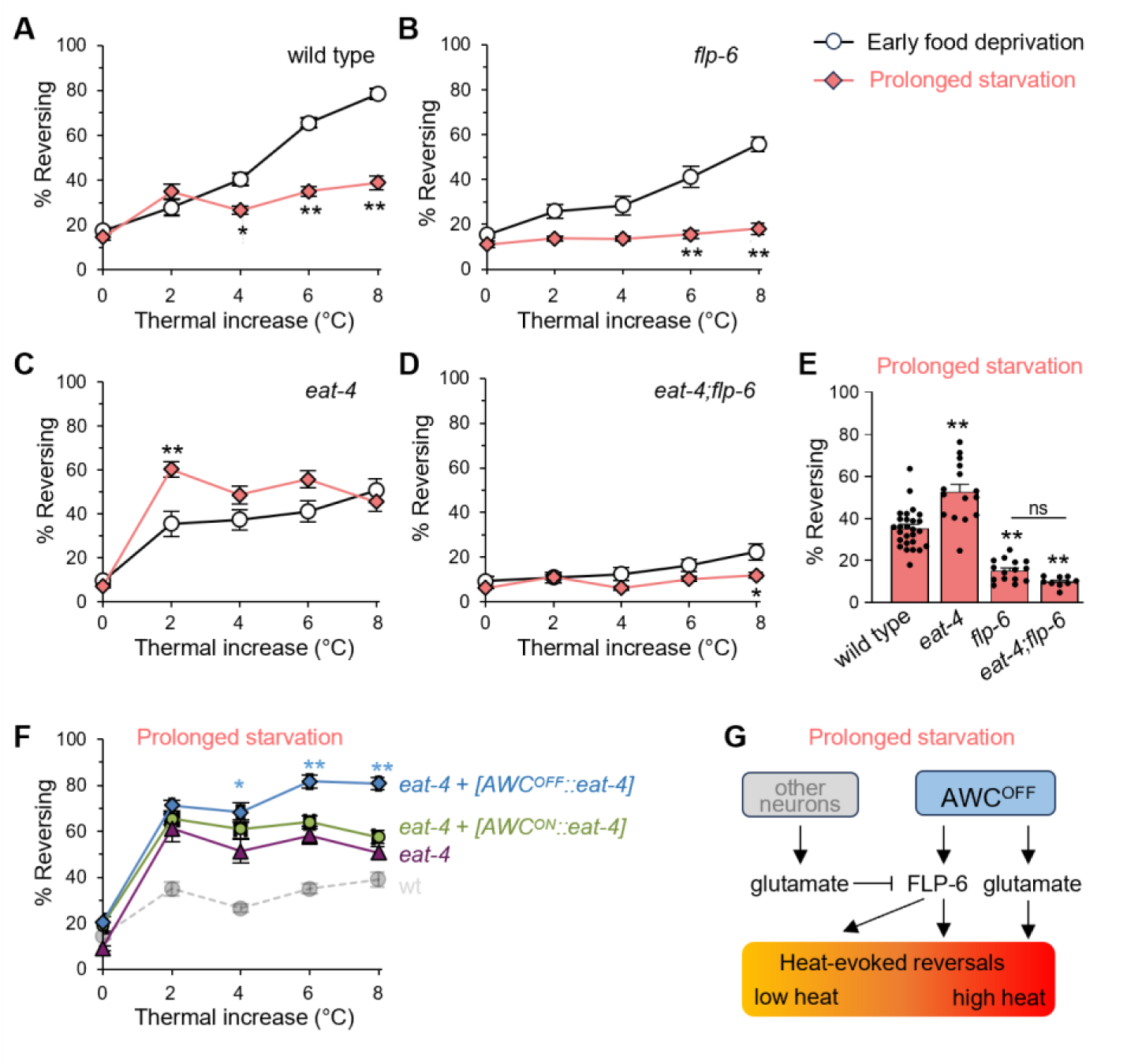
Bidirectional glutamate signaling actions modulate heat-evoked reversals following starvation. **A-D** Comparison of heat-evoked reversals after early food deprivation (off-food 1hr) and prolonged starvation (off-food 6hr) in wild type, *eat-4(ky5)*, *flp-6(ok3056)* and double mutant. **E** Heat-evoked reversal rate averaged over the four heating levels for the prolonged starvation condition. Same data as in panel A-D. **F** Analysis of transgenic animals with AWC subtype-specific rescue of *eat-4* after prolonged starvation (off-food 6hr). Results are presented as average +/- S.E.M. \**\*, p<.01* between the two food-deprivation timepoints (A-E) and versus non-transgenic mutants (F) at respective heat levels, by Bonferroni post-hoc tests. The number of assays (*n*), each scoring at least 50 worms, were: wild type, *n=*27; *eat-4*, *n=*15; *flp-6*, *n=*15; *eat-4+[AWC^OFF^::eat-4]*, *n=*6; *eat-4+[AWC^ON^::eat-4]*, *n=*9. **G** Visual model illustrating the bidirectional effect of glutamatergic signaling from AWC^OFF^ and from unidentified non-AWC neurons (other).

Third, since ASI is required for starvation-evoked plasticity but does not release classical neurotransmitters, we hypothesized that its function is mediated by neuropeptidergic signaling. We therefore screened for defects in starvation-dependent plasticity in mutants for neuropeptide genes known to be expressed in ASI: *ins-4, ins-6, ins-32,* and *nlp-18* [26, 34, 35, 40]*. ins-4* mutants exhibited normal plasticity, with robust responsiveness under early food deprivation and significantly reduced responses after prolonged starvation (Figure 7A). Mutants for *ins-6, ins-32,* and *nlp-18* all showed reduced responsiveness in the early food deprivation condition, but abnormally elevated responses after prolonged starvation (Figure 7A). This phenotype was qualitatively similar to the potentiation phenotype of *eat-4* mutants (Figure 6C). We performed further analyses with *ins-32* and *nlp-18* mutants, since they produced the most significant potentiation over a large range of heat intensities. To determine whether ASI is a relevant source of INS-32 and NLP-18 neuropeptides, we conducted ASI-specific rescue experiments in *ins-32* and *nlp-18* mutants. In *ins-32* mutant background, ASI-specific expression of *ins-32* produced a strong rescue effect, significantly restoring reversal response in early food-deprived animals (Figure 7B) and significantly reducing heat-evoked reversal response under the starvation condition (Figure 7C). In *nlp-18* mutant background, ASI-specific expression of *nlp-18* partially restored normal responsiveness under both the early food-deprivation and the prolonged starvation conditions (Figure 7 D and E). Although they do not exclude the possibility of contributions from other sources, these results suggest that ASI is a functionally relevant source of both INS-32 and NLP-18 to promote heat-evoked reversal upon early food-deprivation and to inhibit such reversals after prolonged starvation, (Figure 7F).

**Figure 7.**
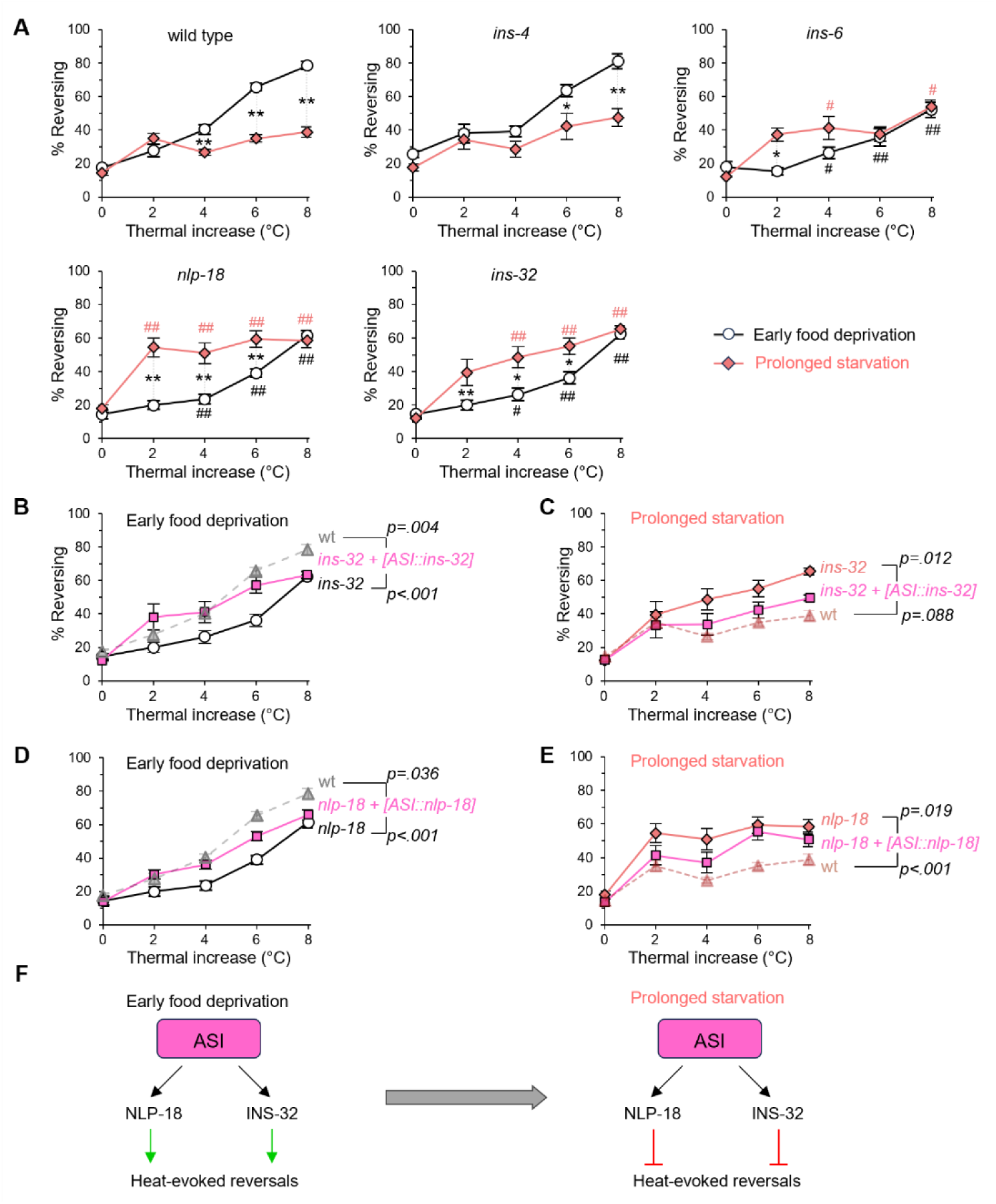
Starvation reconfigures neuropeptidergic signaling by ASI. **A** Comparison of heat-evoked reversals after early food deprivation (off-food 1hr) and prolonged starvation (off-food 6hr) in wild type, *ins-4(ok3534), ins-6(tm2008), nlp-18(ok1557)* and *ins-32(tm6109)*. *, *p<.05,* and \**\*, p<.01* between the two food-deprivation timepoints, by Bonferroni post-hoc tests. #, *p<.05,* and *##, p<.01* versus wild type (N2) at the corresponding timepoint, by Bonferroni post-hoc tests. Data for wild type are the same as the one depicted in Figure 6. **B-E** Impact of transgenic rescue in *ins-32* (B-C) and *nlp-18* (D-E) background. Two-way ANOVA showing no significant heating power x genotype interaction, but a significant genotype main effect, post-hocs tests were conducted on the genotype factor only. Indicated *p* values were corrected with Bonferroni correction. **F** Model of the bidirectional actions and the sources for NLP-18 and INS-32 neuropeptides in the modulation of heat-evoked reversal. Red: reversal-suppressing pathway; Green: reversal-promoting pathway. Additional neuropeptides (such as INS-6) may also be involved, but in the absence of direct evidence for their origin from ASI, they were not included in this scheme.

Collectively, our results (Figure 6 and 7) show that starvation-dependent thermonociceptive plasticity is orchestrated by multiple opposing signaling pathways in which neuropeptides and glutamate signals from different neurons produce context-dependent, bidirectional effects.

## DISCUSSION

Our study reveals that starvation profoundly modulates thermonociceptive behavior in *C. elegans* by engaging distinct neurons, altering neural response polarities, and mobilizing opposing neuromodulatory pathways. This behavioral plasticity, manifesting as a progressive attenuation of heat-evoked reversals under prolonged food deprivation, depends on a shift in the internal state that is not rescued by food-associated olfactory cues. By dissecting the underlying circuits and signaling mechanisms, we uncover two key regulatory nodes involving AWC and ASI neurons, which work in order to integrate thermal and nutritional cues into an adaptive behavioral output, as depicted in the visual model in Figure 8. AWC and ASI, as well as other (unidentified) neurons, use glutamatergic signaling and multiple neuropeptides to orchestrate this plastic response.

**Figure 8.**
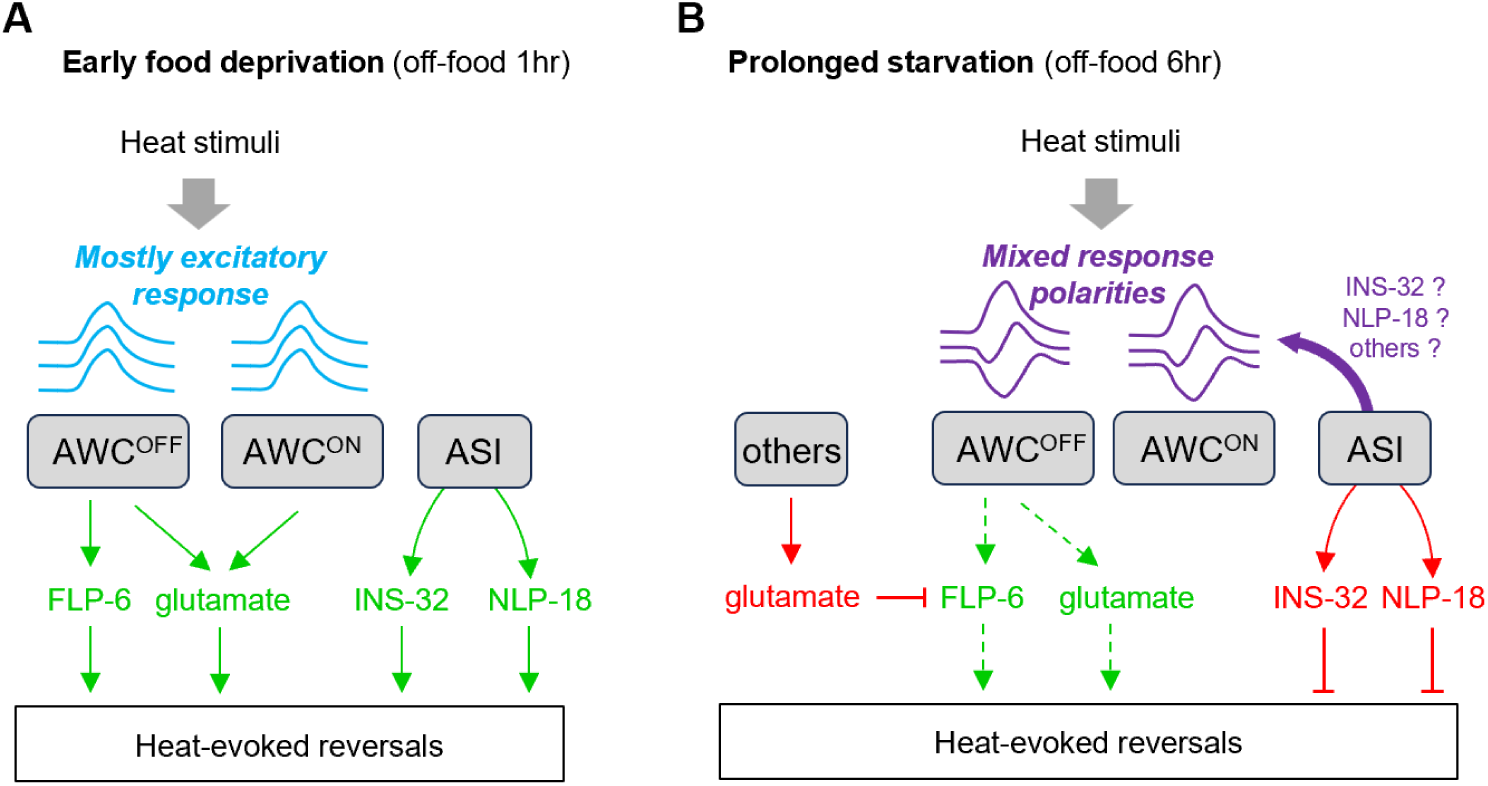
Visual model of the cellular and molecular signaling pathways mediating and modulating heat-evoked reversals according to food deprivation duration. **A** Situation upon early food deprivation (off-food 1hr). The two AWC neurons produce mostly stimulus-locked calcium elevations in response to heat stimuli and make a major contribution to heat-evoked reversals. Glutamate signaling from AWC^ON^ and AWC^OFF^, as well as FLP-6 neuropeptide signaling from AWC^OFF^ mediates heat-evoked reversals. INS-32 and NLP-18 neuropeptide from ASI neurons, as well as potentially from additional unidentified sources (not depicted), also promote heat-evoked reversals. **B** Situation following prolonged starvation (off-food 6hr). Starvation causes several functional changes. First, AWC calcium activity pattern shifts from a mostly excitatory response (calcium elevations) to less predictable response patterns combining calcium elevations (excitation), calcium decreases (inhibition) and variable responses; this effect is mediated by ASI (purple arrow). Second, INS-32 and NLP-18 neuropeptides switch from a reversal-promoting effect to a reversal-suppressing effect. One simple model for INS-32 and NLP-18 action would be that they mediate the ASI effect on AWC activity patterns (purple arrow), but this possibility remains hypothetical. Third, glutamatergic signaling switches from a mostly reversal-promoting effect to a reversal-suppressing effect; this switch is associated with different glutamate sources (AWC and non-AWC neurons, respectively). Of note, residual reversal response upon starvation might be mediated by FLP-6 and glutamate from AWC^OFF^. In summary, our findings highlight a complex reconfiguration of the thermoresponsive circuit following starvation, which includes thermosensory encoding change and complex neuromodulation using bidirectional signaling molecules that produce reversal-promoting (green pathways) or reversal-suppressing effects (red pathways) in a context-dependent manner.

### Starvation regulates thermonociceptive and negative thermotaxis plasticity via at least partly different mechanisms

Previous studies showed that AWC plays an important role in starvation-dependent plasticity in the negative thermotaxis behavior in an innocuous thermal range between 15 and 25°C [26, 33]. Negative thermotaxis involves the detection of thermal changes created by animal movement in spatial thermogradient (0.5°C/cm), the magnitude of the expected thermal changes approximating 0.01°C/s [26]. The starvation impact on negative thermotaxis was shown to (i) involve an up-regulation of AWC cell activity, (ii) rely on INS-1 neuropeptide produced in the intestine and (iii) to occur independently of ASI neurons. In contrast, our study used thermo-nociceptive stimuli, with faster raising thermal slopes (∼0.5-2°C/s, hence 50-200 times faster than those occurring for thermotaxis) and covering noxious temperatures (up to 28°C). Our results indicate that the regulation of thermo-nociceptive response by starvation (i) is linked to a shift in the distribution of AWC activity response polarities from mostly excitatory to a mix of excitatory and inhibitory response, (ii) relies on ASI and specific neuropeptide produced in ASI, and (iii) works independently of INS-1 neuropeptide. Therefore, our study complements our understanding of the modulation of temperature-dependent behavior in *C. elegans* with previously undocumented mechanisms at the circuit, cellular and molecular levels.

### AWC neurons are central drivers of heat-evoked reversal behavior via glutamate and neuropeptides

Our findings confirm that AWC sensory neurons are critical mediators of thermonociceptive behavior. Previous experiments carried out with extreme heating powers (10 °C raise within tens of milliseconds, [21]) had shown differential responsiveness in the two AWCs subtypes and a large behavioral impact was found in mutants affecting AWC asymmetries. The heating intensity range used in the present study (∼0.5 to 2°C per second) triggered a similar withdrawal response, which almost entirely relied on the presence of intact AWC neurons. However, under our experimental conditions, we did not find major differences between AWC^ON^ or AWC^OFF^ subtypes in terms of calcium dynamics, nor did mutations affecting their asymmetry produce radical effects. Therefore, noxious heat-evoked activity in AWC varies widely according to context, which is in line with previous literature [18, 20, 41]. Interestingly, the role of AWC is distinct from that of AFD and FLP neurons, which are canonically linked to thermosensation and nociception [5, 14, 42, 43], but contribute only modestly to heat-evoked behavior in our assay conditions with between 1 and 6 hrs of food deprivation.

We further demonstrate that AWC neurons use both glutamatergic and neuropeptidergic signals to encode heat intensity, with glutamate (via EAT-4) primarily driving responses to stronger stimuli and the FLP-6 neuropeptide acting over a broader thermal range. Functional asymmetry between AWC^ON^ and AWC^OFF^ emerges in our rescue experiments, with AWC^OFF^ more effective in restoring heat-evoked behavior, particularly through FLP-6. This suggests that AWC^OFF^ may have privileged access to neuropeptidergic regulatory machinery or downstream targets, although glutamatergic signaling appears largely redundant across AWC subtypes. Together with previous findings, our results refine our appreciation of how AWC responds to heat and how developmental asymmetries might modulate aversive response to thermal cues over a very broad intensity range.

### Starvation reshapes AWC heat stimulus-locked response from predominantly excitatory to a pattern combining excitatory and inhibitory responses

A key observation of this work is that prolonged starvation transforms heat stimulus-locked AWC calcium responses from intensity-dependent excitatory response patterns to a more heterogeneous response mode. This shift is not merely a quantitative attenuation but a qualitative reorganization of response states, including frequent calcium downregulation or variable traces, which in the same animal can comprise both calcium up- and down-regulation of calcium levels during a single stimulus train. We speculate that this transition from mostly excitatory responses to a mix of excitatory/inhibitory response profiles reflects a fundamental reconfiguration of sensory processing under prolonged nutrient deprivation. In an early food-deprived state, AWC neurons may signal heat intensity in a consistent manner to promote aversion. Under prolonged starvation, however, dampened and variable AWC responses may bias the animal away from escape behaviors, thus prioritizing food-seeking over threat avoidance. This state-dependent reconfiguration may serve as a flexible strategy to balance risk avoidance and metabolic needs, as elaborated below.

### ASI neurons modulate AWC dynamics and promote plasticity via neuropeptides

Our data demonstrate that ASI neurons are dispensable for baseline thermonociceptive responses but essential for starvation-induced behavioral plasticity. Ablation of ASI fully abolished the starvation-dependent attenuation of heat-evoked reversals and prevented the pattern shift in AWC calcium response polarities. These results suggest that ASI gates the modulation of AWC activity based on internal nutritional state. ASI might also modulate the circuit downstream of AWC. Interestingly, in two previous studies, starvation-evoked down-regulation of AWC-dependent thermotactic behavior was intact when ASI was either transiently or chronically inhibited [26, 33]. Therefore, starvation most likely uses a different circuitry to modulate noxious-heat evoked reversal than for thermotaxis modulation.

We further identify two ASI-expressed neuropeptides, NLP-18 and INS-32, as relevant mediators of this plasticity. Mutants for the corresponding pro-neuropeptide genes displayed abnormal behavioral adaptation to starvation, and transgenic expression of either peptide in ASI partially rescues the plasticity defect. ASI was previously shown to mediate starvation-evoked phenotypic plasticity regarding developmental trajectories (via TGFbeta and the insulin-like peptide DAF-28 [44, 45]), food-seeking exploration in starved larvae (via NLP-24 opioid-like neuropeptide [46]), the curvature taken during head-touch evoked escape response (via NLP-18 [47]), and ASH-mediated octanol avoidance response (via multiple NLP neuropeptides [48]). Our findings now add context-dependent aversive thermosensory behavior regulation to the growing list of satiety signaling and stress integration-related functions regulated by ASI neurons. It remains to be determined how ASI integrates systemic cues of metabolic status and how its neuropeptidergic outputs interact with AWC-neural pathway at the synaptic or circuit level to regulate noxious heat avoidance. Another open question is whether ASI action takes place during development (prior to starvation), or more acutely with active signaling after starvation.

### Opposing neuromodulatory influences shape behavioral flexibility in order to balance protection and exploration strategies

The balance between starvation-induced suppression and preservation of thermonociceptive behavior appears to be controlled by a set of signaling molecules, whose effect can be bidirectional, operating in a context-dependent manner. Upon early food deprivation, neuropeptides, including INS-32 released from ASI and NLP-18 from an unidentified source, as well as glutamatergic signals from AWC neurons promote thermonociceptive behaviors (Figure 8A). Upon prolonged starvation, the same signaling molecules can produce the opposite effect and suppress this behavior (Figure 8B). For NLP-18 and glutamate the differential effect is associated with a potentially different source, e.g., non-AWC neurons for glutamate. For INS-32, our cell-specific rescue experiment suggests that ASI is a functionally relevant source in the two contexts. However, we cannot rule out the implication of additional neurons, nor an overexpression effect. While additional studies will be needed to decipher the context-dependent mode of action of each of these signaling molecules, our findings suggest a regulatory architecture in which bidirectional neuromodulatory signals integrate metabolic context with sensory drive to determine behavioral priorities. Rather than globally silencing the thermonociceptive pathway, starvation might well reconfigure it through a layered combination of neuronal state shifts, circuit-level inhibition, and peptide-based tuning. This complexity may afford the animal greater flexibility in selecting context-appropriate responses under competing environmental and internal pressures. From an ecological and evolutionary perspective, the downregulation of nociceptive responses during prolonged starvation likely reflects a strategic shift in behavioral priorities. In the face of acute noxious stimuli, animals typically favor rapid escape to minimize injury, a protective response. However, under prolonged food deprivation, such protection may come at the cost of missed opportunities for locating food. In this context, a heightened aversive response could lead to excessive avoidance of mildly threatening environments that may nevertheless contain nutritional resources. In addition, because worm exploration strategies rely on modulating locomotion speed, the duration of forward locomotion bouts and the frequency of re-orientation events, often associated with reversals, the down-regulation of reversal response is directly relevant to the rate of animal dispersion in the environment [49, 50]. By reducing the sensitivity or reliability of nociceptive pathways, particularly through modulating AWC output, *C. elegans* may adopt a more exploratory behavioral mode, tolerating greater risk to increase the chances of finding food. This trade-off between protection and exploration mirrors similar state-dependent shifts observed in other species [51, 52] suggesting a conserved strategy wherein internal metabolic needs dynamically reshape threat evaluation and behavioral responses.

## METHODS

### *C. elegans* maintenance and strains list

*C. elegans* strains were maintained according to standard techniques on nematode growth medium (NGM) agar plates seeded with OP50. Animal synchronization was made by treating gravid adults with standard hypochlorite-based procedure. File S1 includes a list of strains used in the present study. The lines with genetical neuron ablation were previously described [8, 22, 53].

### Cloning and transgenesis

Promoter-containing Entry plasmids (Multi-site Gateway slot 1) were constructed by PCR using N2 genomic DNA as template and primers flanked by attB4 and attB1r recombination sites; the PCR product being cloned into pDONR-P4-P1R vector (Invitrogen) by BP recombination. Coding sequence-containing Entry plasmids (Multi-site Gateway slot 2) were constructed by PCR using N2 cDNA as template and primers flanked by attB1 and attB2 recombination sites; the PCR product being cloned into pDONR_221 vector (Invitrogen) by BP recombination. Expression plasmids for transgenesis were created through LR recombination reactions (Gateway LR Clonase, Invitrogen) as per the manufacturer’s instructions. DNA prepared with a GenElute HP Plasmid miniprep kit (Sigma) was microinjected in the gonad to generate transgenic lines according to a standard protocol [54]. Rescue constructs included SL2::mCherry transgenes, the expression pattern of which was verified using epifluorescence microscopy, as previously described [55]. File S1 includes a list of plasmids and primers used in this study.

### Behavioral assays

#### Preparation of worms

All the behavioral experiments were performed in young adult animals cultivated at 20°C. For the Fed condition, 80 animals were transferred to a NGM plate fully covered with OP50 bacterial lawn 18 h before the experiment. For starved condition, fed animals were washed 3 times in 1.5 mL collection tubes with M9 (pre-equilibrated at the growth temperature of worms) to remove OP50. During washes, worms were left to settle to the bottom of the tubes by gravity. About 80 animals were then plated on unseeded NGM plates with a drop of M9 and left to air dry for 5 min. Duration of starvation was considered from the time of the start of the wash. During starvation, animals were kept in the incubator at 20°C. Food-odor experiments were performed by adding OP50 bacteria on the inward side of the petri dish lid during the starvation period.

#### Heat-stimulation protocol

Heat stimulation was delivered using previously described INFERNO system [31]. The stimulation program consisted of a baseline period of 40 s without any heat stimulation (to determine spontaneous reversal rate), 4 s with 100 W heating (1 IR lamp turned on), 20 s of interstimulus interval (ISI), 4 s with 200 W heating (2 lamps turned on), 20 s of ISI, 4 s with 300 W heating (3 lamps turned on), 20 s of ISI and 4 s with 400 W heating (4 lamps turned on) as previously described. Thermal increases were measured with Type K thermal probes linked to Data Logger (Pico Technology). For these four heating power levels, the peak temperatures reached at the surface of the plate were ∼22, ∼24, ∼26 and ∼28°C, respectively, corresponding to thermal slope approximating 0.5, 1, 1.5 and 2 °C/s, respectively.

#### Movie recording and analysis

During the stimulation program, worm plates were filmed using a DMK 33U×250 camera and movies acquired with the IC capture software (The Imaging Source), at a 1600×1800 pixel resolution, at 8 frames per second, and the resulting .AVI file was encoded as Y800 8-bit monochrome. Behavioral recordings were analyzed using the Multi-Worm Tracker 1.3.0 (MWT) [56]. A custom Python script was used to flag reversal events and report the frame during which they occurred. The movie time course was separated into 4-s bins and the fraction of animals reversing in each bin was extracted as the primary output for subsequent analyses.

### Calcium imaging

#### Worm preparation and thermal stimulation

The imaging was performed 1 h and 6 h after food deprivation. Animals cultivated at 20°C were glued, maintained at 20°C for 90 s of baseline imaging acquisition and subjected to 30-s heat stimulation at 22°C, 24°C, 26°C and 28°C using CherryTemp microfluidic system by Cherry Biotech, with 60-s ISI. The system works by shifting between two fluidic loops that each pass through a thermalization block located in the vicinity of the specimen. It takes between 4 and 6 s to achieve 80% of the thermal shift between baseline (20°) and the higher temperature target.

#### Data acquisition and trace processing

Calcium imaging was performed using a Leica DMI6000B inverted epifluorescence microscope equipped with a HCX PL Fluotar L40x/0.60 CORR dry objective, a Leica DFC360FX CCD camera, an EL6000 light source, and equipped with fast filter wheels for FRET imaging (excitation filter: 427 nm (BP 427/10); emission filters 472 (BP 472/30) and 542 nm (BP 542/27) [14]. Calcium levels in AWC^ON^ and AWC^OFF^ neurons were monitored using cameleon YC2.3 sensor. Imaging YFP/CFP emission ratio (R) of the YC2.3 cameleon indicator was used to analyze relative changes in response to short-lasting stimuli. Baseline drift was estimated by linear interpolation between consecutive baseline periods (average over 10 s prior to each stimulus) and subtracted from the raw ratiometric trace (to obtain ΔR values), effectively removing slow, non-physiological signal components (due to differential photobleaching of the two fluorophores) while preserving rapid stimulus-evoked calcium responses. Baseline-corrected ΔR traces were used for visualization and comparative analyses, including heat maps of response amplitude and directions.

#### Manual trace categorization

Trace categorization as *calcium up*, *calcium down* or *variable* was done manually (not blind to the condition). First, the polarity of calcium signal change was assessed on a trial-by trial-basis. For each trial, we considered the direction of the first visible peak following thermal stimulation and labeled it as either *calcium up trial, calcium down trial,* or *undefined polarity trial* (when no obvious direction could be delineated due to limited signal change or to low signal-to-noise ratio). A visible peak was defined as a peak whose magnitude was at least twice as large as the signal variation (noise) seen in the immediately preceding 10-s baseline period and whose duration was at least 2 frames. Second, traces were categorized as *calcium up* (with three or more *calcium up* trails), as *calcium down traces* (with three or more *calcium down* trails) and as *variable* (other cases). Globally, this manual categorization used arbitrary signal-to-noise cutoff and criteria to qualitatively assess traces and sort them for visualization.

#### Peak intensities histograms

To complement the manual trace categorization with a more quantitative approach, we computed the response magnitude in each by averaging the ΔR signal in the 15-s post stimulus (this ΔR signal already subtracts the local baseline signal). The distributions of response magnitudes (which could be either positive for excitatory responses or negative for inhibitory responses) were presented in histograms.

### Statistical analyses

We used Jamovi (Version 2.6.44) to conduct Kruskal-Wallis non-parametric tests (to compare calcium peak distribution differences) and ANOVAs, followed by Bonferroni post hoc tests (to compare behavioral scores). Multiple-comparison corrections took account of repeated comparisons to the same controls. Some analyses are reported across multiple figure panels, in which case the same control data was plotted several times, with indication in the respective figure legends. Replicate numbers (*n*) are defined in the Figures. These *n* were determined in agreement with previous studies using similar measures. No *a priori* power analyses were performed. Results of statistical tests, including *p*-values and effect sizes, are included in File S2.

## ACKNOWLEDGEMENTS

We are grateful to Lisa Schild and Laurence Bulliard for expert technical support, to Aurore Jordan for the cloning of AWC-specific promoter plasmids and for the participation in calcium trace acquisition. Some strains were provided by the CGC, which is funded by NIH Office of Research Infrastructure Programs (P40 OD010440). Some strains were provided by NBRP, which is funded by the Japanese government. The study was supported by the Swiss National Science Foundation (BSSGI0_155764, PP00P3_150681, and 310030_197607 to DAG).

## DATA AVAILIBILITY STATEMENT

The data associated with this article are included in the manuscript, the figure and supplements (with raw data in File S2).

## MATERIAL AVAILIBILITY STATEMENT

The strains and plasmids generated during the present study should be requested to the corresponding author.

## Figure and Figure Legends

**Figure 2-supplement 1.**
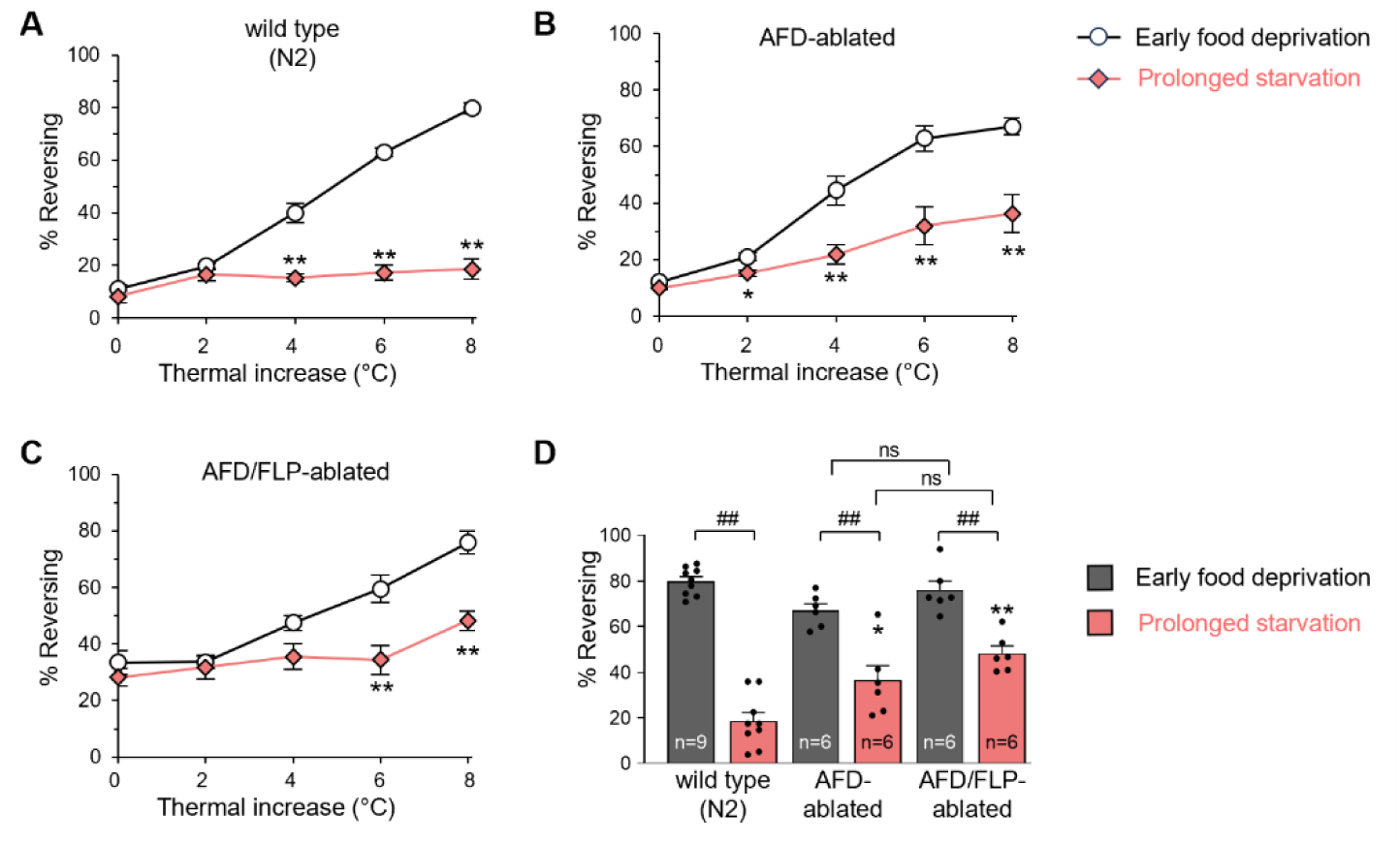
Heat-evoked reversal upon early food deprivation and starvation-dependent plasticity are largely intact in AFD and FLP-ablated animals. Impact of the genetic ablation of candidate thermo-responsive neurons on thermonociceptive response and starvation-dependent plasticity. **A-C** Comparison of heat-evoked reversals after early food deprivation (off-food 1hr) and starvation (off-food 6 hr) in wild type and in transgenic animals with caspase-mediated ablation of indicated neurons. Results are presented as average +/- S.E.M. *, *p<.05,* and \**\*, p<.01* versus corresponding heat level in the early food deprivation condition, by Bonferroni post-hoc tests. **D** Comparison for the highest heating level (thermal increase = 8 °C) across the three genotypes presented in panel A to C. Bars as average, dots as individual assay scores, and error bars as S.E.M. *##, p<.01* between early food deprivation and prolonged starvation for each genotype; \*\**, p<.01* versus wild type (N2) at the corresponding timepoint, by Bonferroni post-hoc tests. The total number of assays (n) analyzed per condition, each scoring at least 50 worms, are indicated in panel D. The wild type (N2) dataset is the same as in Figure 2D. Data reported in the two figures were acquired in parallel and multiple comparison corrections were made in a single analysis.

**Figure 3-Supplement 1.**
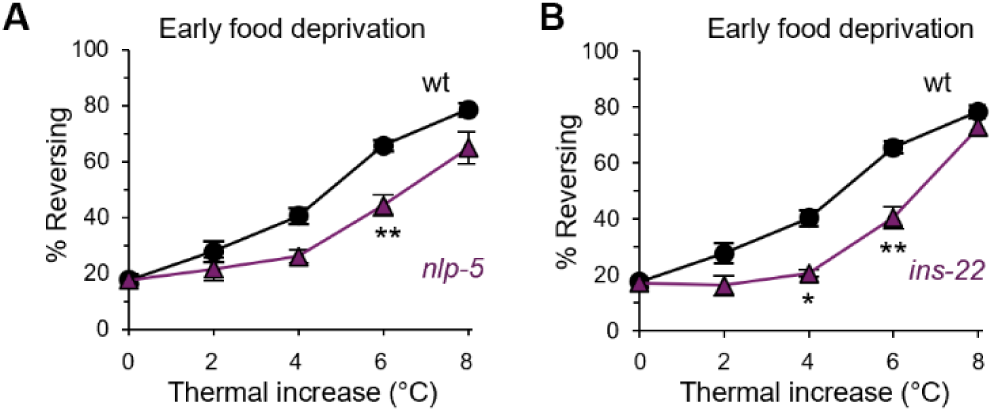
Alteration of heat-evoked reversals in *nlp-5* and *ins-22* mutants. **A-B** Comparison of heat-evoked reversals after early food deprivation (off-food 1hr) in wild type, *nlp-5(ok1981)* and *ins-22(ok3616)*. Results are presented as average +/- S.E.M. *, *p<.05* and \**\*, p<.01* versus wild type by Bonferroni post-hoc tests. The number of assays (*n*), each scoring at least 50 worms, were: wild type, *n=*27; *nlp-5*, *n=*6; *ins-22*, *n=*6. A single wt control dataset is reported across panels in this Figure and in Figure 3.

**Figure 4-supplement 1.**
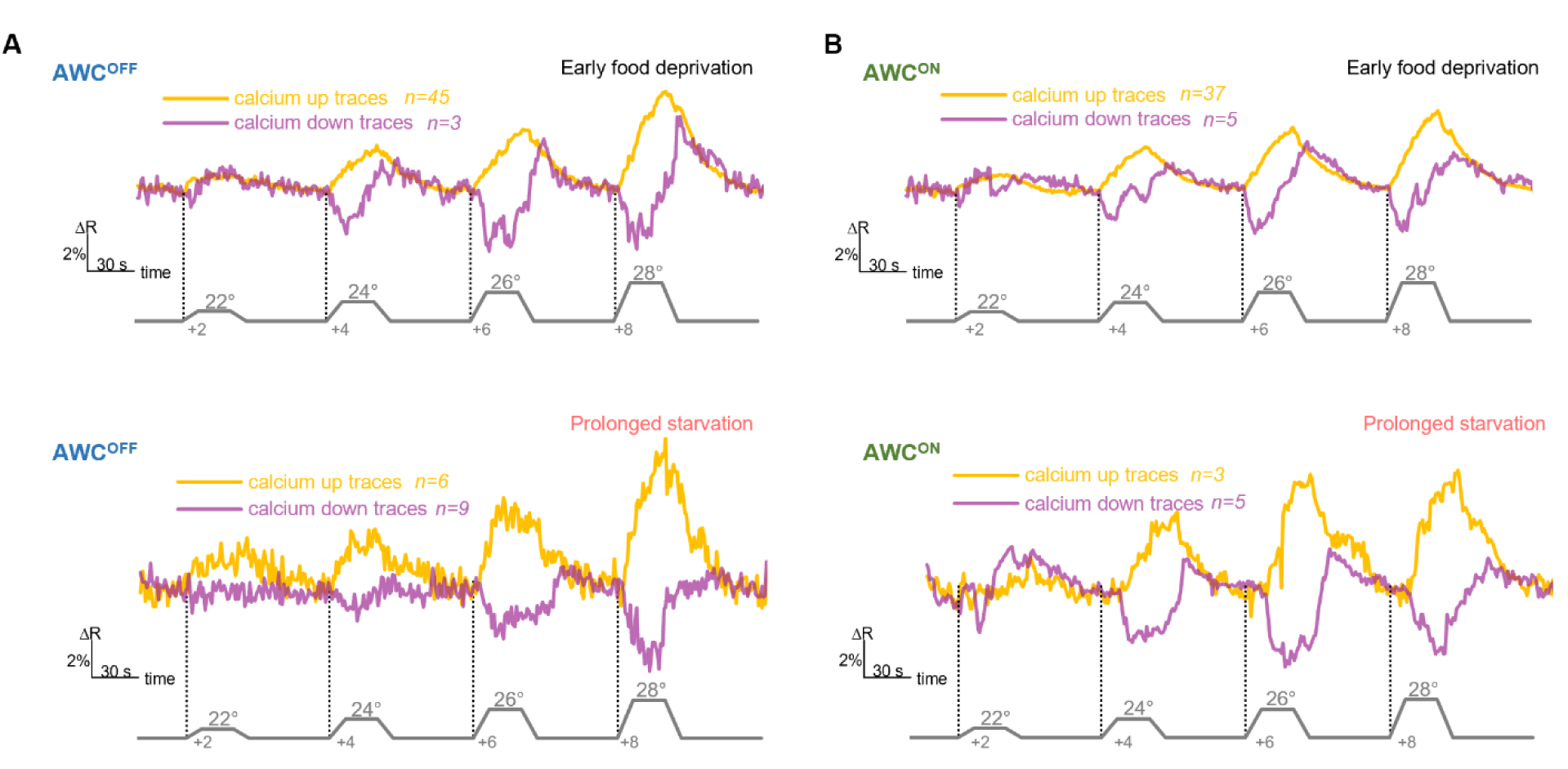
Similar kinetics of calcium up and calcium down responses under the early food deprivation and the prolonged starvation condition. **A-B** From the same dataset as in Figure 4 A-B, average traces showing the dynamics of *calcium up* (excitatory) or *calcium down* (inhibitory) response types with similar amplitude and globally comparable shapes.

**Figure 6-Supplement 1.**
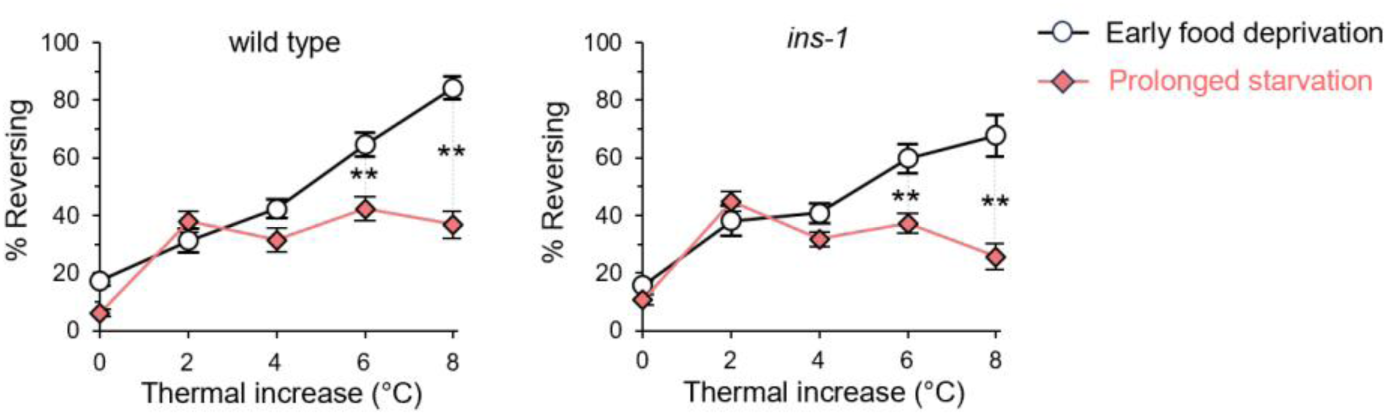
Intact starvation-induced thermonociceptive plasticity in *ins-1* mutants. Comparison of heat-evoked reversals after early food deprivation (off-food 1hr) and prolonged starvation (off-food 6hr) in wild type (N2), and *ins-1(nj32).* *, *p<.05,* and \**\*, p<.01* between the two food deprivation timepoints, by Bonferroni post-hoc tests. #, *p<.05,* and *##, p<.01* versus wild type (N2) at the corresponding timepoint, by Bonferroni post-hoc tests.

## REFERENCES

1. Ezcurra, M., et al., Food sensitizes C. elegans avoidance behaviours through acute dopamine signalling. EMBO J, 2011. 30(6): p. 1110–1122.

2. Ryu, L., et al., Feeding state regulates pheromone-mediated avoidance behavior via the insulin signaling pathway in Caenorhabditis elegans. Embo j, 2018. 37(15).

3. Burrell, B.D., Comparative biology of pain: What invertebrates can tell us about how nociception works. J Neurophysiol, 2017. 117(4): p. 1461–1473.

4. Byrne Rodgers, J. and W.S. Ryu, Targeted thermal stimulation and high-content phenotyping reveal that the C. elegans escape response integrates current behavioral state and past experience. PLOS ONE, 2020. 15(3): p. e0229399.

5. Liu, S., E. Schulze, and R. Baumeister, Temperature- and Touch-Sensitive Neurons Couple CNG and TRPV Channel Activities to Control Heat Avoidance in Caenorhabditis elegans. PLoS One, 2012. 7(3): p. e32360.

6. Jordan, A. and D.A. Glauser, Distinct clusters of human pain gene orthologs in Caenorhabditis elegans regulate thermo-nociceptive sensitivity and plasticity. Genetics, 2023. 224(1).

7. Rudgalvyte, M., et al., Antagonist actions of CMK-1/CaMKI and TAX-6/calcineurin along the C. elegans thermal avoidance circuit orchestrate adaptation of nociceptive response to repeated stimuli. Elife, 2025. 14.

8. Glauser, D.A., et al., Heat avoidance is regulated by transient receptor potential (TRP) channels and a neuropeptide signaling pathway in Caenorhabditis elegans. Genetics, 2011. 188(1): p. 91–103.

9. Schild, L.C., et al., The balance between cytoplasmic and nuclear CaM kinase-1 signaling controls the operating range of noxious heat avoidance. Neuron, 2014. 84(5): p. 983–96.

10. Wittenburg, N. and R. Baumeister, Thermal avoidance in Caenorhabditis elegans: an approach to the study of nociception. Proc Natl Acad Sci U S A, 1999. 96(18): p. 10477–82.

11. Mohammadi, A., et al., Behavioral response of Caenorhabditis elegans to localized thermal stimuli. BMC Neurosci, 2013. 14(1): p. 66.

12. Glauser, D.A. and M.B. Goodman, Molecules empowering animals to sense and respond to temperature in changing environments. Curr Opin Neurobiol, 2016. 41: p. 92–98.

13. Goodman, M.B. and P. Sengupta, The extraordinary AFD thermosensor of C. elegans. Pflugers Arch, 2018. 470(5): p. 839–849.

14. Saro, G., et al., Specific Ion Channels Control Sensory Gain, Sensitivity, and Kinetics in a Tonic Thermonociceptor. Cell Rep, 2020. 30(2): p. 397–408 e4.

15. Thapliyal, S., I. Beets, and D.A. Glauser, Multisite regulation integrates multimodal context in sensory circuits to control persistent behavioral states in C. elegans. Nat Commun, 2023. 14(1): p. 3052.

16. Bargmann, C.I., Chemosensation in C. elegans. WormBook, 2006: p. 1–29.

17. Colbert, H.A. and C.I. Bargmann, Odorant-specific adaptation pathways generate olfactory plasticity in C. elegans. Neuron, 1995. 14(4): p. 803–12.

18. Kuhara, A., et al., Temperature sensing by an olfactory neuron in a circuit controlling behavior of C. elegans. Science, 2008. 320(5877): p. 803-7.

19. Zaslaver, A., et al., Hierarchical sparse coding in the sensory system of Caenorhabditis elegans. Proc Natl Acad Sci U S A, 2015. 112(4): p. 1185–9.

20. Biron, D., et al., An olfactory neuron responds stochastically to temperature and modulates Caenorhabditis elegans thermotactic behavior. Proc Natl Acad Sci U S A, 2008. 105(31): p. 11002–7.

21. Kotera, I., et al., Pan-neuronal screening in Caenorhabditis elegans reveals asymmetric dynamics of AWC neurons is critical for thermal avoidance behavior. Elife, 2016. 5.

22. Beverly, M., S. Anbil, and P. Sengupta, Degeneracy and neuromodulation among thermosensory neurons contribute to robust thermosensory behaviors in Caenorhabditis elegans. J Neurosci, 2011. 31(32): p. 11718–27.

23. Harris, G., et al., Dissecting the serotonergic food signal stimulating sensory-mediated aversive behavior in C. elegans. PLoS One, 2011. 6(7): p. e21897.

24. Mills, H., et al., Monoamines and neuropeptides interact to inhibit aversive behaviour in Caenorhabditis elegans. Embo j, 2012. 31(3): p. 667–78.

25. Oranth, A., et al., Food Sensation Modulates Locomotion by Dopamine and Neuropeptide Signaling in a Distributed Neuronal Network. Neuron, 2018. 100(6): p. 1414–1428.e10.

26. Takeishi, A., et al., Feeding state functionally reconfigures a sensory circuit to drive thermosensory behavioral plasticity. Elife, 2020. 9.

27. Rengarajan, S., et al., Feeding state sculpts a circuit for sensory valence in Caenorhabditis elegans. 2019. 116(5): p. 1776-1781.

28. Banerjee, N., et al., Differential processing of a chemosensory cue across life stages sharing the same valence state in Caenorhabditis elegans. 2023. 120(19): p. e2218023120.

29. Ghosh, D.D., et al., Neural Architecture of Hunger-Dependent Multisensory Decision Making in C. elegans. Neuron, 2016. 92(5): p. 1049–1062.

30. Ezcurra, M., et al., Neuropeptidergic Signaling and Active Feeding State Inhibit Nociception in Caenorhabditis elegans. J Neurosci, 2016. 36(11): p. 3157–69.

31. Lia, A.S. and D.A. Glauser, A system for the high-throughput analysis of acute thermal avoidance and adaptation in C. elegans. J Biol Methods, 2020. 7(1): p. e129.

32. Ippolito, D., S. Thapliyal, and D.A. Glauser, Ca2+/CaM binding to CaMKI promotes IMA-3 importin binding and nuclear translocation in sensory neurons to control behavioral adaptation. eLife, 2021. 10: p. e71443.

33. Yeon, J., A. Takeishi, and P. Sengupta, Chronic vs acute manipulations reveal degeneracy in a thermosensory neuron network. MicroPubl Biol, 2021. 2021.

34. Ghaddar, A., et al., Whole-body gene expression atlas of an adult metazoan. Sci Adv, 2023. 9(25): p. eadg0506.

35. Taylor, S.R., et al., Molecular topography of an entire nervous system. Cell, 2021. 184(16): p. 4329–4347.e23.

36. Ghosh, R., et al., Multiparameter behavioral profiling reveals distinct thermal response regimes in Caenorhabditis elegans. BMC Biol, 2012. 10: p. 85.

37. Lesch, B.J., et al., Transcriptional regulation and stabilization of left-right neuronal identity in C. elegans. Genes Dev, 2009. 23(3): p. 345–58.

38. Troemel, E.R., A. Sagasti, and C.I. Bargmann, Lateral signaling mediated by axon contact and calcium entry regulates asymmetric odorant receptor expression in C. elegans. Cell, 1999. 99(4): p. 387–98.

39. Gallagher, T., et al., ASI regulates satiety quiescence in C. elegans. J Neurosci, 2013. 33(23): p. 9716–24.

40. Chen, Y. and L.R. Baugh, Ins-4 and daf-28 function redundantly to regulate C. elegans L1 arrest. Developmental Biology, 2014. 394(2): p. 314–326.

41. Flavell, S.W., D.M. Raizen, and Y.J. You, Behavioral States. Genetics, 2020. 216(2): p. 315–332.

42. Kimura, K.D., et al., The C. elegans Thermosensory Neuron AFD Responds to Warming. Current Biology, 2004. 14(14): p. 1291–1295.

43. Chatzigeorgiou, M. and William R. Schafer, Lateral Facilitation between Primary Mechanosensory Neurons Controls Nose Touch Perception in C. elegans. Neuron, 2011. 70(2): p. 299–309.

44. Makino, M., et al., Regulation of Satiety Quiescence by Neuropeptide Signaling in Caenorhabditis elegans. Front Neurosci, 2021. 15: p. 678590.

45. Li, W., S.G. Kennedy, and G. Ruvkun, daf-28 encodes a C. elegans insulin superfamily member that is regulated by environmental cues and acts in the DAF-2 signaling pathway. Genes Dev, 2003. 17(7): p. 844–58.

46. Park, J.Y., et al., A novel functional cross-interaction between opioid and pheromone signaling may be involved in stress avoidance in Caenorhabditis elegans. Sci Rep, 2020. 10(1): p. 7524.

47. Chen, L., et al., CKR-1 orchestrates two motor states from a single motoneuron in C. elegans. iScience, 2024. 27(4): p. 109390.

48. Hapiak, V., et al., Neuropeptides amplify and focus the monoaminergic inhibition of nociception in Caenorhabditis elegans. J Neurosci, 2013. 33(35): p. 14107–16.

49. Glauser, D.A., How and why Caenorhabditis elegans uses distinct escape and avoidance regimes to minimize exposure to noxious heat. Worm, 2013. 2(4): p. e27285.

50. Schild, L.C. and D.A. Glauser, Dynamic switching between escape and avoidance regimes reduces Caenorhabditis elegans exposure to noxious heat. Nat Commun, 2013. 4.

51. Lee, Y.-F., Y.-M. Kuo, and W.-C. Chu, Energy state affects exploratory behavior of tree sparrows in a group context under differential food-patch distributions. Frontiers in Zoology, 2016. 13(1): p. 48.

52. Grunwald Kadow, I.C., State-dependent plasticity of innate behavior in fruit flies. Current Opinion in Neurobiology, 2019. 54: p. 60–65.

53. Russell, J., et al., Humidity sensation requires both mechanosensory and thermosensory pathways in Caenorhabditis elegans. Proc Natl Acad Sci U S A, 2014. 111(22): p. 8269–74.

54. Evans, T.C., (ed), Transformation and microinjection. WormBook, 2006.

55. Marques, F., et al., Tissue-specific isoforms of the single C. elegans Ryanodine receptor gene unc-68 control specific functions. PLoS Genet, 2020. 16(10): p. e1009102.

56. Swierczek, N.A., et al., High-throughput behavioral analysis in C. elegans. Nat Methods, 2011. 8(7): p. 592–8.

