## Supplementary material for "Starvation transforms signal encoding in *C. elegans* thermoresponsive neurons and suppresses heat avoidance via bidirectional glutamatergic and peptidergic signaling": File S1

**Promoter plasmids (multisitegateway slot 1)**

Entry plasmids containing specific promoters were constructed by PCR from N2 genomic DNA with primers flanked with attB4 and attB1r recombination sites and cloned into pDONR-P4-P1R vector (Invitrogen) by BP recombination. Primer sequences were the following:

dg950 [slot1 Entry str-2p]

attB4str-2_F: ggggacaactttgtatagaaaagttgATTGACGTAGATGTGGGCTTGAAGTA

attB1rstr-2_R: ggggactgcttttttgtacaaacttgGATCACGAGTATTCGGACAAAAAAGTCT

dg949 [slot1 Entry srsx-3p]

attB4srsx-3_F: ggggacaactttgtatagaaaagttgATGCAGTTGGCTCAAGGTTTGTTATG

attB1rsrsx-3_R: ggggactgcttttttgtacaaacttgCCACAGTCGAGCTgATTCCGAA

dg833 [slot1 Entry gpa-4p]

attB4gpa-4_F: ggggacaactttgtatagaaaagttgATTGCGACTTTCGATACGTAGG

attB1rgpa-4_R: ggggactgcttttttgtacaaacttgTTGTTGAAAAGTGTTCACAAAATG

**Coding sequence plasmids (multisitegateway slot 2)**

Entry plasmids containing specific coding DNA sequences (cds) or genomic sequence (gs) were constructed by PCR from N2 cDNA or N2 genomic DNA with primers flanked with attB1 and attB2 recombination sites and cloned into pDONR_221 vector (Invitrogen) by BP recombination. Primer sequences were the following:

dg1014 [slot2 Entry flp-6cds]

attB1flp-6_F: ggggacaagtttgtacaaaaaagcaggctTAATGAACTCTCGTGGGTTGATTTTGA

attB2flp-6_R: ggggaccactttgtacaagaaagctgggtCTTATCGTCCGAATCTCATGTATGCT

dg830 [slot2 Entry ins-32cds]

attB1ins-32_F: ggggacaagtttgtacaaaaaagcaggctTAATGACCTCGATTCTGTTGATCC

attB2ins-32_R: ggggaccactttgtacaagaaagctgggtCTCATTCATCAGGGCACATTATTGA

dg831 [slot2 Entry nlp-18cds]

attB1nlp-18_F: ggggacaagtttgtacaaaaaagcaggctTAATGAACGCCAACGTCTACTCGA

attB2nlp-18_R: ggggaccactttgtacaagaaagctgggtCTTAGGCGAACGATGAGAATTGTCC

dg718 [slot2 YC2.3 cameleon] previously described in (Saro et al. 2020)

**3’ UTR and tagging plasmids (multi-site gateway slot3)**

mg277 [SL2::mCherry] was created previously (Schild *et al*, 2014). mg211 [EntrySlot3unc-54UTR] (aka pMH473) was a gift from Marc Hammarlund.

**Expression plasmids used for transgenesis**

dg785 [str-2p::QF::unc-54UTR] was created through a LR recombination reaction between dg950, dg240, mg211, and pDEST-R4-P3.

dg783 [srsx-3p::QF::unc-54UTR] was created through a LR recombination reaction between dg949, dg240, mg211, and pDEST-R4-P3.

dg845 [gpa-4p::QF::unc-54UTR] was created through a LR recombination reaction between dg833, dg240, mg211, and pDEST-R4-P3.

dg839 [QUASp::flp-6cds::SL2::mCherry] was created through a LR recombination reaction between dg229, dg827, mg277, and pDEST-R4-P3.

dg838 [QUASp::eat-4cds::SL2::mCherry] was created through a LR recombination reaction between dg229, dg713, mg277, and pDEST-R4-P3.

dg842 [QUASp::ins-32cds::SL2::mCherry] was created through a LR recombination reaction between dg229, dg830, mg277, and pDEST-R4-P3.

dg843 [QUASp::nlp-18cds::SL2::mCherry] was created through a LR recombination reaction between dg229, dg831, mg277, and pDEST-R4-P3.

**Strain list:**

| **Strain Name** | **Genotype** | **Comments** |
| --- | --- | --- |
| N2 | Wild type | Wild type (WT) |
| PY7502 | *ceh-36delp::TU#813; ceh-36delp::TU#814; srtx-1p::gfp; Coelomycte::dsRED* | AWC ablation (Gift from Piali Sengupta).  Fig. 2B, D |
| PY7505 | *gpa-4p::TU318; gcy-27p::TU814; gcy-27p::GFP coelp::dsRED* | ASI ablation (Gift from Piali Sengupta).  Fig. 2C, D |
| GN112 | *pgIs2 [gcy-8p::TU#813 + gcy-8p::TU#814 + unc-122p::GFP + gcy-8p::mCherry + gcy-8p::GFP + ttx-3p::GFP]* | AFD ablation (gift from Miriam B. Goodman). Supplementary Fig. 2B, D |
| JPS481 | *vxEx280 [sto-5p::ICE + gcy-8::ICE + myo-2p::mCherry]* | AFD and FLP ablation. Obtained from CGC  Supplementary Fig. 2C, D |
| MT6308 | *eat-4(ky5) III* | Obtained from CGC.  Fig. 3A, 3B, 6C, 6D |
| DAG1156-1157 | *eat-4(ky5) III; domEx1156-1157[str-2p::QF 20 ng/ul, QUASp::eat-4CDS::SL2mcherry 20 ng/ul, unc122p::RFP 20 ng/ul]* | *eat-4* rescue with str*-2* promoter driving eat*-4* coding sequence (cds).  Fig. 3B, 6D |
| DAG1231-1232 | *eat-4(ky5) III; domEx1231-1232[srsx-3p::QF 60 ng/ul, QUASp::eat-4CDS::SL2mcherry 20 ng/ul, unc122p::RFP 20 ng/ul]* | *eat-4* rescue with srsx*-3* promoter driving eat*-4* coding sequence (cds).  Fig. 3B, 6D |
| VC2324 | *flp-6(ok3056) V* | Obtained from CGC.  Fig. 3C, 3D, 6B |
| DAG1802-1803 | *flp-6(ok3056) V; domEx1802-1803[str-2p::QF 20 ng/ul, QUASp::flp-6_SL2mcherry 20 ng/ul, unc122p::GFP 20 ng/ul]* | *flp-6* rescue with *str-2* promoter driving *flp-6* coding sequence (cds).  Fig. 6D |
| DAG1904-1905 | *flp-6(ok3056) V; domEx1904-1905[srsx-3p::QF 60 ng/ul, QUASp::flp-6_SL2mcherry 20 ng/ul, unc122p::GFP 20 ng/ul]* | *flp-6* rescue with *srsx-3* promoter driving *flp-6* coding sequence (cds).  Fig. 6D |
| RB1609 | *nlp-5(ok1981) II.* | Obtained from CGC.  Supplementary Fig. 3.1A |
| RB2594 | *ins-22(ok3616) III.* | Obtained from CGC.  Supplementary Fig. 3.1B |
| VC390 | *nsy-1(ok593) II.* | Obtained from CGC.  Supplementary Fig. 3.2A |
| tm3080 | *nsy-7 (tm3080)* | Obtained from NBRP Japan.  Supplementary Fig. 3.2B |
| DAG1417 | *domEx1417[srsx-3p::QF 60 ng/ul, QUASp::YC2.3CDS::unc-54UTR 20 ng/ul, unc122p::RFP 20 ng/ul]* | YC2.3 Cameleon in AWC^OFF^  Fig. 4 |
| DAG1148 | *domEx1418[str-2p::QF 20 ng/ul, QUASp::YC2.3CDS::unc-54UTR 20 ng/ul, unc122p::RFP 20 ng/ul]* | YC2.3 Cameleon in AWC^ON^  Fig. 4 |
| DAG1901 | *domEx1417[srsx-3p::QF, QUASp::YC2.3CDS::unc-54UTR, unc122p::RFP]; [gpa-4p::TU318, gcy-27p::TU814, gcy-27p::GFP, coelp::dsRED]* | YC2.3 Cameleon in AWC^OFF^ in ASI neurons ablated background  Fig. 5 |
| DAG1902 | *domEx1148[str-2p::QF::UTR54, QUASp::YC2.3::UTR54, unc122p::RFP]; [gpa-4p::TU318, gcy-27p::TU814, gcy-27p::GFP, coelp::dsRED]* | YC2.3 Cameleon in AWC^ON^ in ASI neurons ablated background  Fig. 5 |
| RB2544 | *ins-4(ok3534) II.* | Obtained from CGC.  Fig. 7A |
| tm2008 | *ins-6(tm2008)* | Obtained from NBRP Japan.  Fig. 7A |
| RB1372 | *nlp-18(ok1557) II.* | Obtained from CGC.  Fig. 7A |
| tm6109 | *ins-32(tm6109)* | Obtained from NBRP Japan.  Fig. 7A |
| DAG1892-DAG1902 | *ins-32(tm6109); domEx1892, 1906[gpa-4p::QF 30 ng/ul, QUASp::ins-32CDS::SL2mcherry 30 ng/ul, unc122p::GFP 20 ng/ul]* | *ins-32* rescue with *gpa-4* promoter driving *ins-32* coding sequence (cds).  Fig. 7B-C |
| DAG1237-DAG1238 | *nlp-18(ok1557) II; domEx1237-1238[gpa-4p::QF 30 ng/ul, QUASp::nlp-18CDS::SL2mcherry 30 ng/ul, unc122p::GFP 20 ng/ul]* | *nlp-18* rescue with *gpa-4* promoter driving *nlp-18* coding sequence (cds).  Fig. 7D-E |
